# N-terminal phosphorylation regulates the activity of Glycogen Synthase Kinase 3 from *Plasmodium falciparum*

**DOI:** 10.1101/2021.06.18.448949

**Authors:** Samuel Pazicky, Arne Alder, Haydyn Mertens, Dmitri Svergun, Tim Gilberger, Christian Löw

## Abstract

As the decline of malaria cases stalled over the last five years, novel targets in *Plasmodium falciparum* are necessary for the development of new drugs. Glycogen Synthase Kinase (PfGSK3) has been identified as a potential target, since its selective inhibitors were shown to disrupt the parasite’s life cycle. Here, we show that PfGSK3 exhibits autophosphorylation, leading to an extensive phosphorylation both *in vitro* and in the parasite. In the uncanonical N-terminal region of the parasite enzyme, we identified several autophosphorylation sites that regulate the activity of PfGSK3. By combining molecular modeling with experimental small-angle X-ray scattering data, we show that increased PfGSK3 activity is promoted by conformational changes in the PfGSK3 N-terminus, triggered by N-terminal phosphorylation. Our work provides novel insights into the structure and regulation of the malarial PfGSK3.

## Introduction

*Plasmodium falciparum* is an intracellular eukaryotic parasite that causes the most severe form of malaria in humans, infecting over 200 million people every year (1). The emergence of malaria parasites resistant to all currently used antimalarial drugs imposes a serious threat to public health (2). Identification of new drug targets and drugs is therefore crucial for future handling of the malaria pandemics. Kinases comprise up to 10% of all drug targets (3) and a number of potent inhibitors were discovered that are directed against *P. falciparum* kinases, such as glycogen synthase kinase 3 (PfGSK3) (4). Human GSK3ß plays a crucial role in a number of signaling processes, regulating glycogen metabolism (5–7), cell cycle and growth (8–10), translation (7), embryonic development (9, 11) or differentiation of neurons (12–16). Consequently, the inhibitors of GSK3 are investigated as potential treatment options for neurodegenerative and psychiatric diseases (17–23) and its structure, catalytic mechanism and regulation are well studied (24–26). GSK3β is thought to be constitutively activated by autophosphorylation at Y216 (27–29) and inactivated by phosphorylation of N-terminal S9 mediated by several protein kinases, such as PKA and PKB (6, 30). The inactivation is achieved as the phosphorylated S9 enables binding of GSK3 N-terminus to a pocket for a priming phosphorylation in the substrate-binding region and thus reduces the binding affinity for substrates (24, 25).

In *P. falciparum*, two genes are encoding GSK3 (PF3D7_0312400 and PF3D7_1316000) and these enzymes are much less understood in their physiological role and in the molecular mechanism. GSK3ß (PF3D7_0312400, further as PfGSK3) is the only isoform that is expressed in blood stages (31) of the parasite and is involved in the regulation of red blood cell invasion (32). The invasion process is an essential, parasite-driven event that relies on the orchestrated interplay of multiple protein-protein interactions (33). It requires the apical membrane antigen 1 (AMA1), a type I transmembrane protein located in the micronemes of the parasite and translocated on its surface before the host cell entry (32, 34–39). The N-terminal ectodomain of AMA1 interacts with the rhoptry neck (RON) protein complex of the parasite, that is inserted into the membrane of the red blood cell, forming the basis of the so called “tight junction” between the membrane of the invading parasite and its target cell (40– 44). The short C-terminus of AMA1 locates inside the parasite and thus, is amenable to regulation by phosphorylation (45, 46). In a two-step phosphorylation event, protein kinase A (PfPKA) first phosphorylates S610 of AMA1 (45, 47, 48), which in turn enables subsequent phosphorylation of AMA1 residue T613 by PfGSK3 (32). Thus, PfGSK3 serves as a secondary kinase, although both phosphorylation sites are equally important for AMA1 function and *P. falciparum* invasion (32). PfGSK3 inhibitors, built on a thieno[2,3-b]pyridine or benzofuran scaffold, have been developed that display selective antiplasmodial activity with IC_50_ values in the low micromolar to sub-micromolar range (49–51) and one of these inhibitors has been shown to abrogate the red blood cell invasion (32). Further improvements of these inhibitors could be guided by a three-dimensional structure of PfGSK3 that has, however, not been determined so far.

Based on sequence homology across GSK3 proteins, PfGSK3 consists of two conserved structural domains: a β-sheet-rich lobe with an ATP binding pocket and an α-helical lobe with the activation loop and a substrate binding site. These domains are typically preceded by an N-terminal domain that is predicted to be disordered (25). In PfGSK3, the important residues in the ATP binding pocket and in the activation loop remain conserved, however, its N-terminus displays an unique extension and its function remains unknown (52).

To analyze this in more detail, we optimized the purification protocol of PfGSK3 that enabled us to perform thorough structural and functional characterization of the protein *in vitro*. We show evidence that PfGSK3 exhibits autophosphorylation of the N-terminal and activation loop residues *in vitro* and validated these phosphorylation sites *in vivo*. Phosphorylation of the N-terminal residues induces conformational changes of the protein that lead to an increase in the enzymatic activity of PfGSK3, providing a new regulatory mechanism.

## Results

### High yield expression and purification of enzymatically active PfGSK3 from *E. coli*

To obtain recombinant PfGSK3 in high purity and homogeneity that allow functional and structural analysis, we extended the previously established PfGSK3 expression and purification protocols (32, 49, 50, 53). In short, N-terminally hexahistidine-tagged PfGSK3 (Fig. 1A) was separated from the bacterial cell lysate by immobilized metal affinity chromatography (IMAC, Fig. 1B) and subsequent size exclusion chromatography (SEC, Fig. 1C). We deliberately excluded a negative IMAC step as PfGSK3 becomes sensitive to NiNTA beads upon the removal of the histidine tag and has a high tendency to precipitate. Instead, to remove the impurities, we included an additional ion exchange chromatography (IEC) step (Fig. 1D). This yields the desired level of purity of PfGSK3 as judged by SDS-PAGE, but the protein elutes in several partially overlapping peaks with increasing concentration of sodium chloride. Such heterogeneity of PfGSK3 might be caused by different posttranslational modifications. To assess the quality of the purified protein and to characterize it further, we applied several biophysical methods. Dynamic light scattering data confirm high homogeneity of the sample with a calculated molecular weight of 52 kDa, in agreement with the expected mass for a monomer of recombinant PfGSK3 (52.05 kDa, Fig. S2A). The secondary structure content derived from circular dichroism data is 22% α helix, 26% β sheet, 24% turns and 28% disorder. This corresponds to a typical domain organization of homologous GSK3 proteins (such as human GSK3β) with an N-terminal β-sheet-rich domain and an α-helical C-terminal domain that are flanked by disordered N-terminal and C-terminal regions (Fig. S1A, Fig. 2A) (24, 54). To verify that the protein retains its three-dimensional structure and kinase properties, we tested whether its thermal stability changes in the presence of ATP and ATP analogs. Indeed, the stability of PfGSK3 markedly increased in a concentration dependent manner from 45°C to over 50°C with increasing concentration of dATP or ATP (Fig. 2B and Fig. S2B). Moreover, the protein could also be stabilized by several non-hydrolysable ATP analogs (Fig. S2C). Additionally, we observed a concentration-dependent increase of the thermal stability of PfGSK3 with increasing concentrations of sulfate ions (Fig. S2D-E). Sulfate ions are bound in the activation loop of several human GSK3 structures (55–61), suggesting that the structural elements in this region are also conserved. Finally, we measured the PfGSK3 activity with a luciferase assay, using a peptide derived from human glycogen synthase 1 (GS-1) as a substrate, resulting in a catalytic rate constant of *k*_*cat*_ = 9.5 s^-1^, which is in a similar range as previously reported (between 0.7 and 3.7 s^-1^) for human GSK3 (Fig. 2C, Fig. S2F) (54). Thus, purified PfGSK3 is folded, stable, enzymatically active and retains the structure that is expected to be similar to human GSK3.

**Fig. 1.**
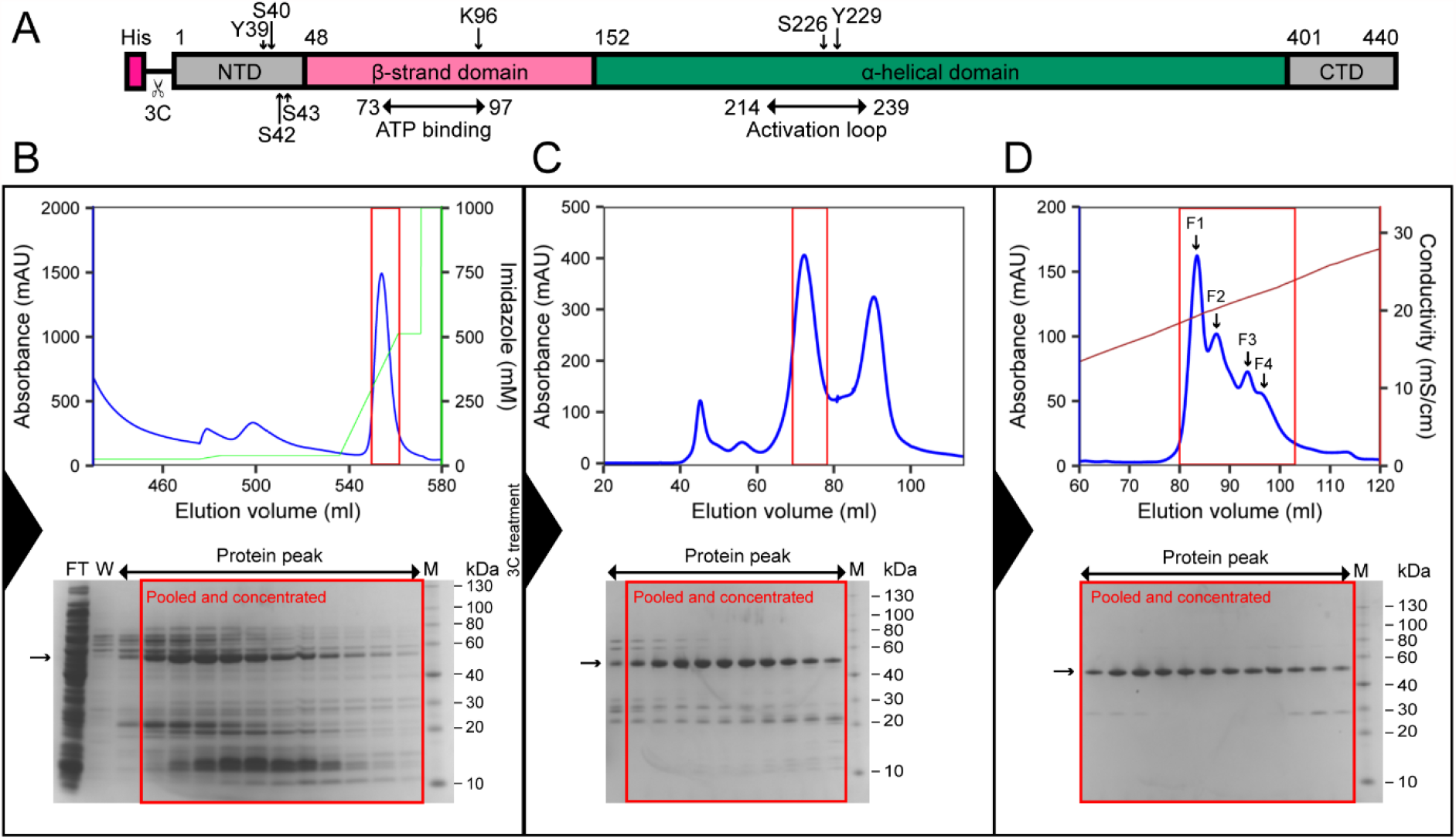
Expression and purification of PfGSK3. (A) Construct of PfGSK3 used for expression. The construct consists of the full-length sequence of PfGSK3 with an N-terminal His-tag and a 3C cleavage site. The domain organization and phosphorylation sites are marked. NTD is the N-terminal domain, CTD is the C-terminal domain, 3C is the 3C protease cleavage cite. (B) First PfGSK3 purification step: imidazole gradient elution profile from the His-Trap column. (C) Elution profile from the Superdex 200 size exclusion column. (D) NaCl gradient elution profile from Resource Q ion exchange column. The peaks of the ion exchange elution represent fractions that are phosphorylated to different extent, increasing from F1 to F4. The blue curves in the chromatograms show UV absorbance, the green curve shows imidazole concentration and the brown curve shows the conductivity. The peaks in red squares in chromatograms correspond to the red squares in corresponding SDS-PAGE gels.

**Fig. 2.**
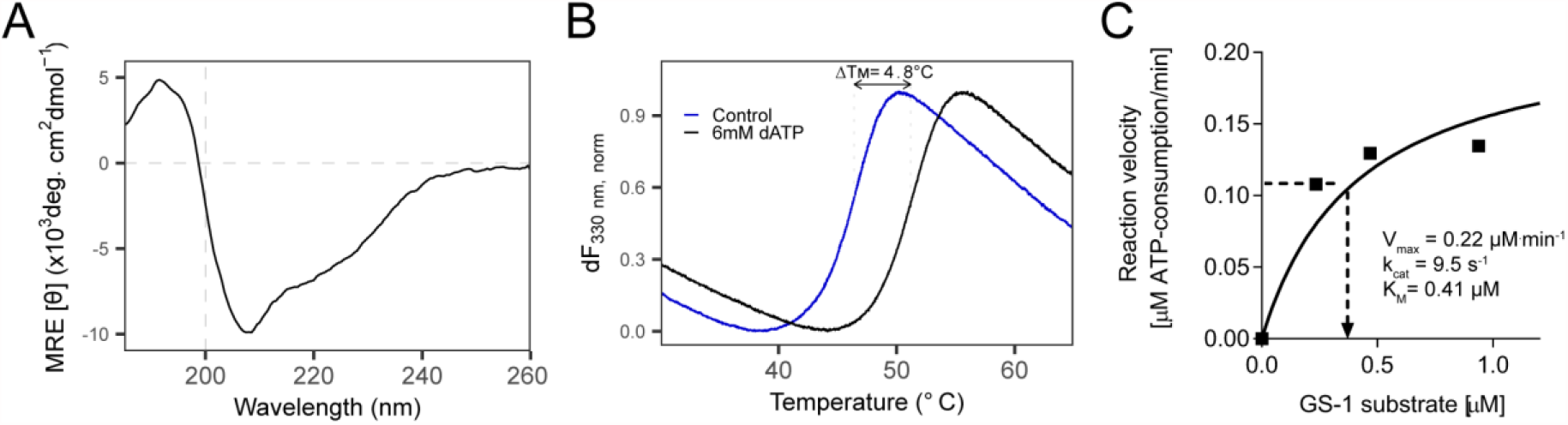
Characterization of PfGSK3. (A) Circular dichroism shows secondary structure composition similar to what is expected for a GSK3 protein, with 22% α helix, 26% β sheet, 24% turns and 29% disordered content. The circular dichroism was measured 10× and the data were averaged, buffer subtracted, and analyzed by DichroWeb. (B) Thermal unfolding profiles of PfGSK3 in presence or absence of 6mM dATP. dATP stabilizes PfGSK3, suggesting that it binds in the ATP binding pocket. (C) Catalytic activity of GSK3 follows Michaelis-Menten kinetics. Saturation curve shows the reaction velocity (as calculated in FigS2F) plotted against different substrate concentrations. Michaelis-Menten constant (K_m_), maximal reaction velocity (V_max_) and turnover number (k_cat_) were calculated in GraphPad Prism using nonlinear regression. GS-1 = substrate peptide of the human glycogen synthase.

### Divalent ions of heavy metals induce reversible formation of PfGSK3 oligomers

The observation that PfGSK3 after affinity tag cleavage has a strong tendency to precipitate upon interaction with NiNTA beads prompted us to investigate the impact of various heavy metals on PfGSK3. To this end, we measured the thermal unfolding profiles of the protein in the presence of different metal ions. Under standard conditions, PfGSK3 displays a clear transition midpoint at 45°C. However, in the presence of divalent ions, such as zinc, cobalt, nickel and copper cations, the typical unfolding transition profiles completely disappear, indicating structural changes (Fig. S3A). We selected zinc to investigate further the effect of divalent heavy metal ions on PfGSK3 because it showed the most pronounced effect on PfGSK3 unfolding. Interestingly, the secondary structure content measured by circular dichroism did not notably change upon addition of zinc, demonstrating that the secondary structural elements of the protein remain folded (Fig. S3C). However, analytical size exclusion chromatography revealed that the zinc cations induce the formation of high molecular weight (high-MW) protein species that elute earlier from the column than the metal-free protein (Fig. 3A) and this effect is dependent on the concentration of the zinc cations (Fig. S3B). Indeed, in a cross-linking experiment with glutaraldehyde, the high-MW species appear predominantly in the presence of zinc (Fig. S3D). The high-MW species can revert to monomers by the addition of the divalent metal chelator EDTA (Fig. 3A). To study the impact of heavy metals on the PfGSK3 function, we measured its activity in presence of zinc at different concentrations. The metal cations cause a dramatic decrease in the PfGSK3 activity (Fig. 3B). However, the activity is completely restored after EDTA treatment. Finally, we used small-angle X-ray scattering (SAXS, Fig. 3C and Fig. S3E) and negative stain EM (Fig. S3F) to characterize the high-MW species. The distance distribution derived from the SAXS data demonstrates high anisometry of the particles (peaking at 20 nm but with *D*_*max*_=90 nm). In agreement, the cross-linked high-MW species visualized by negative stain EM vary in their shape and size (Fig. S3F). In summary, heavy metal ions, such as zinc cations, induce a reversible formation of large PfGSK3 particles. While the protein maintains its secondary structure, it completely loses its enzymatic activity. Importantly, the process is reversible and the restored PfGSK3 monomers fully regain their kinase activity.

**Fig. 3.**
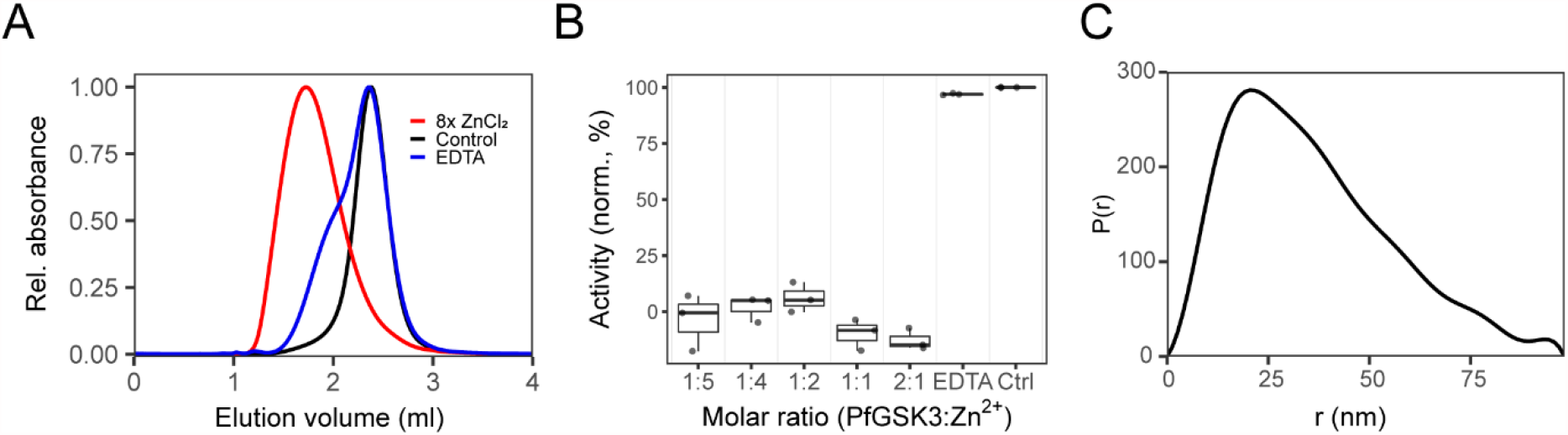
Heavy metal ions inhibit PfGSK3 activity. (A) Analytical size exclusion chromatography profiles on a Superose 6 column of PfGSK3 apoprotein (black), after addition of 8× molar excess of zinc chloride (red) and after subsequent addition of EDTA (blue). The analysis shows that zinc ions induce the formation of high-MW PfGSK3 species. The formation is reversible because addition of EDTA shifts the elution profiles towards higher elution volumes. (B) The activity of PfGSK3 is strongly reduced in the presence of zinc ions, whereas the protein regains the activity after the addition of the metal chelator EDTA. (C) The distance distribution of the PfGSK3 high-MW species induced by zinc ions derived from SAXS data shows high heterogeneity in the sample, peaking at 20 nm but with *D*_*max*_=89 nm.

### PfGSK3 exhibits autophosphorylation

PfGSK3 elutes from the IEX chromatography column in at least four different species with distinct elution volumes that are of comparable purity (Fig. 1D). Such behavior could be explained by a heterogeneous phosphorylation pattern. To investigate this possibility, we separately analyzed the four top peak fractions (F1-F4) from the IEX elution profile of the wild type PfGSK3 preparation (Fig. 1D). SDS-PAGE stained with the Pro-Q Diamond stain, which specifically binds to phosphorylated proteins, shows that the amount of total phosphorylation increases towards later elution volumes (Fig. S4A). Mass spectrometry analysis revealed that the protein is partially phosphorylated at multiple sites. We identified a number of phosphorylated residues at the N-terminus (residues 13-62, see Fig. S4B). Additionally, several residues in the activation loop, primarily S226 and Y229, but also S228 and S232 were found to be phosphorylated (Fig. S4B). The phosphorylation of the activation loop residues and two N-terminal serine residues (S31, S62) is in agreement with the previously identified phosphorylation sites in *P. falciparum* blood-stage parasites (37–39, 62–65), while the phosphorylation of other N-terminal residues, predominantly Y39, S40, S42 and S43, has not been detected in parasite lysate-based phospho-fingerprinting. As the *E. coli* expression system only possesses a limited capacity to phosphorylate proteins (66–68), we tested the ability of PfGSK3 to exhibit autophosphorylation by incubating recombinant protein with ATP/MgCl_2_ and subsequent analysis of the ion exchange chromatography profiles. Indeed, the treatment with ATP/MgCl_2_ led to a drop in the fluorescence signal from the earlier eluting peaks and gain in the fluorescence signal from later eluting peaks, indicating that PfGSK3 is able to exhibit autophosphorylation *in vitro* (Fig. 4A). In addition, we employed the luciferase activity assay to examine PfGSK3 activity in the absence of a substrate peptide. The assay showed the consumption of ATP even in the absence of the substrate peptide that was dependent on the concentration of PfGSK3, confirming that PfGSK3 exhibits autophosphorylation (Fig. 4B).

**Fig. 4.**
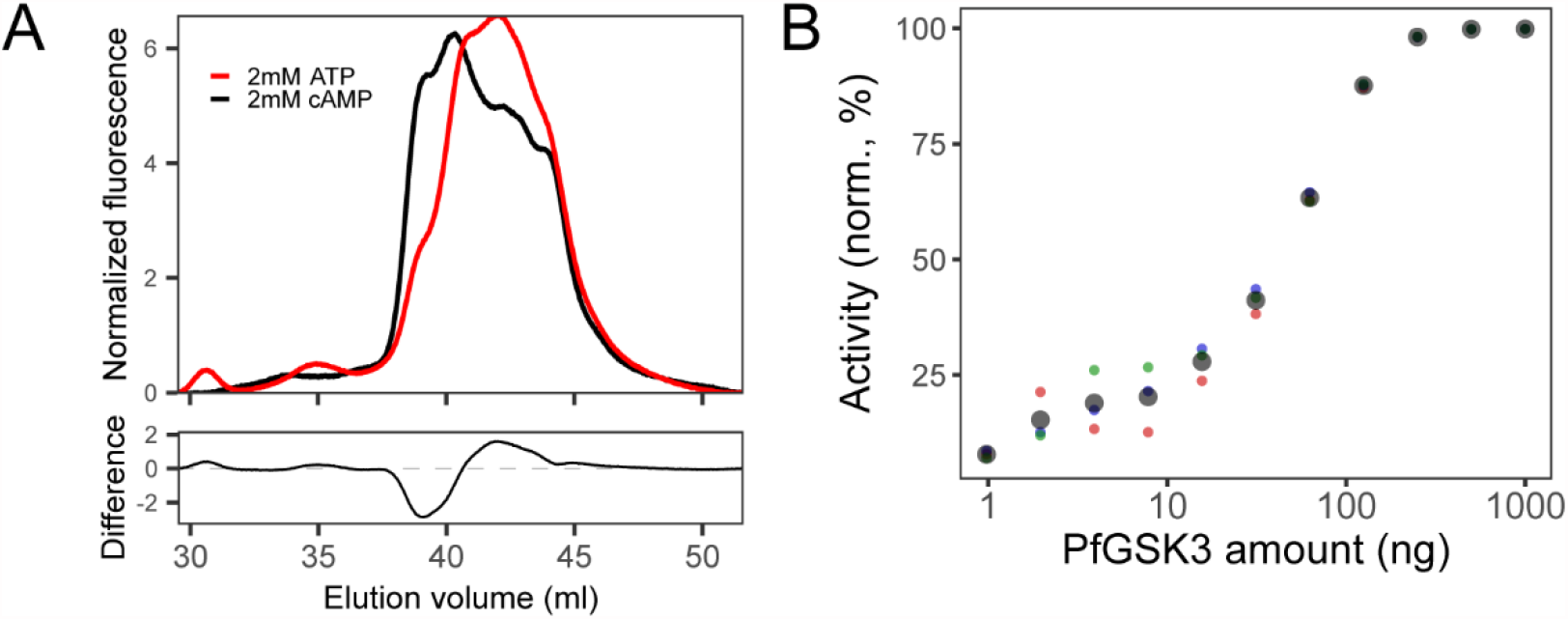
PfGSK3 exhibits autophosphorylation. (A) IEX elution chromatograms (upper plot) of PfGSK3 after incubation with ATP (red) or cAMP as negative control (black) in the presence of magnesium ions suggest that PfGSK3 exhibits autophosphorylation. The difference between both chromatograms (bottom plot) shows an increase of later-eluting species after ATP treatment, indicating higher amount of phosphorylation. (B) The ATP consumption measured by the luminescence assay in the absence of the substrate is dependent on the enzyme concentration, which is indicative of the autophosphorylation exhibited by PfGSK3.

### The N-terminus of endogenous PfGSK3 is phosphorylated

Having observed autophosphorylation of PfGSK3 *in vitro*, we also investigated the phosphorylation state of PfGSK3 expressed in the parasite. To this end, we made use of a *P. falciparum* cell line endogenously expressing C-terminally GFP-tagged GSK3 (GSK3-GFP, Fig. S5). The parasites were harvested at the schizont stage and GSK3-GFP was purified using GFP Trap agarose beads (Fig. 5A) and subjected to mass spectrometry analysis. The identified peptides exhibited phosphorylation both at the N-terminus and in the activation loop, similarly to recombinantly expressed PfGSK3 (Fig. 5B). The generally low Mascot Delta Score (69) did not allow us to confidently pinpoint the exact phosphorylation sites, but confirmed that a significant portion of the residues on the peptide derived from the unstructured N-terminus were phosphorylated (average 28%, residues 39-43) compared to the activation loop peptides (average 5%, residues 226-233). These data attest that N-terminal phosphorylation of PfGSK3 also occurs *in vivo*.

**Fig. 5.**
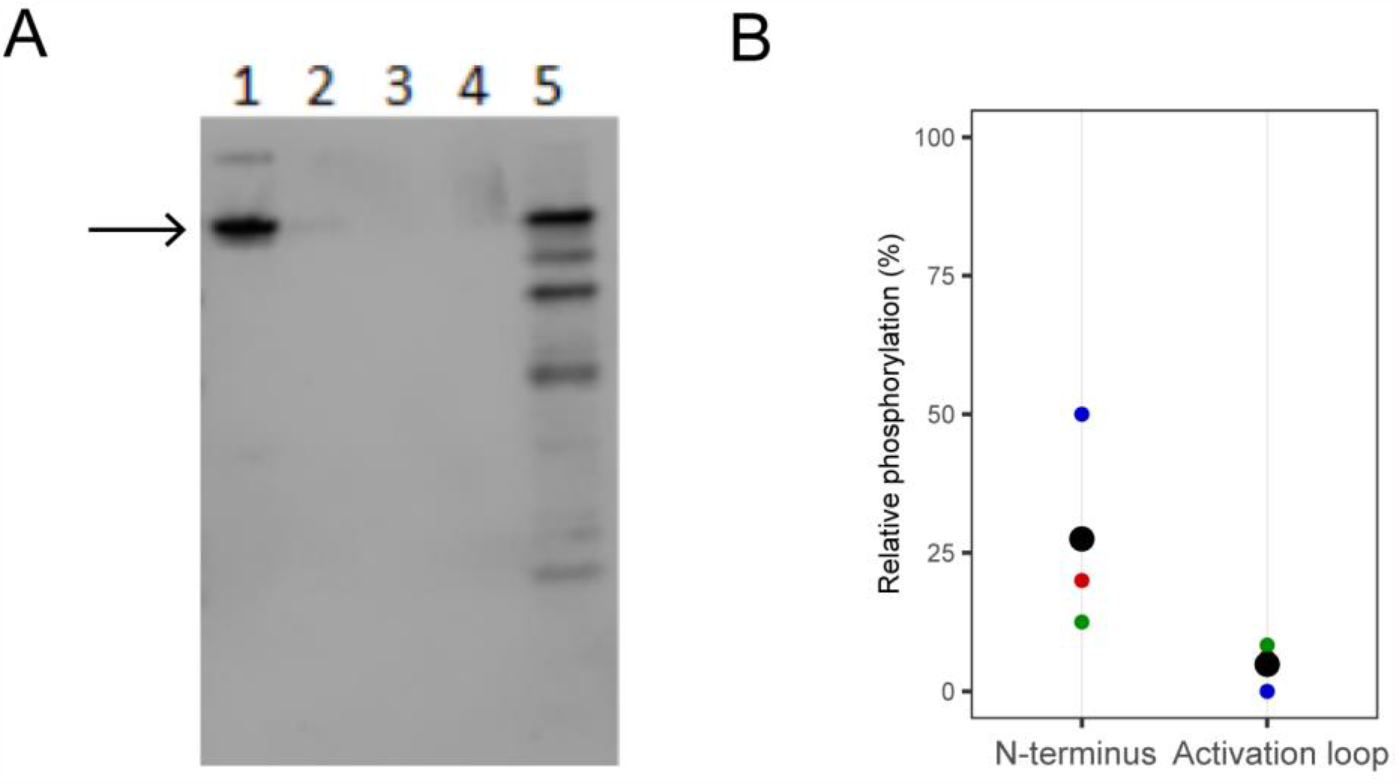
N-terminus of PfGSK3 is phosphorylated *in vivo*. (A) Western blot with anti-GFP antibodies shows the purification of PfGSK3-GFP from the plasmodium lysate (1), flow-through (2), two wash steps (3-4) and elution fraction (5). (B) The mass spectrometry analysis revealed that the N-terminal residues contain higher relative amount of phosphorylation (27%) compared to the activation loop (5%). The colored points are ratios from individual mass spectrometry experiment replicates and the large black points are their averages.

### N-terminal domain is indispensable for PfGSK3 stability

To investigate the role of phosphorylation in PfGSK3 further, we attempted to express inactive mutants to prevent autophosphorylation. Therefore, we changed the essential residue in the ATP binding pocket K96 or residues from the activation loop S226 and Y229 to alanine. The small-scale expression screen showed that the inactive mutants of PfGSK3 did not express in a soluble form but instead formed inclusion bodies (Fig. 6A). We then isolated the inclusion bodies of the PfGSK3^K96A^ and PfGSK3^S226A/Y229A^ mutants and examined their phosphorylation states by mass spectrometry. These mutants display the full length sequence but are devoid of any phosphorylation (K96A) or exhibit markedly reduced phosphorylation (S226A/Y229A) compared to the wild-type protein (Fig. 6C). This indicates that these mutations either impair PfGSK3 function or PfGSK3 phosphorylation is crucial for folding and solubility. The latter is further supported by a co-expression experiment of wild-type PfGSK3 with the dephosphorylating lambda phosphatase, which results in insoluble material only (Fig. S6C). As we found the N-terminal extension of PfGSK3 to carry several phosphorylation sites, we cloned a series of N-terminally truncated protein constructs (Fig. S1A). None of these constructs expressed in a soluble form (Fig. 6B) and mass spectrometry confirmed a complete lack of phosphorylation in the PfGSK3 activation loop (ΔN64 in Fig. 6C). Expression tests at different temperatures indicate that not only the aforementioned mutants and constructs, but also wild-type PfGSK3 is less expressed and mainly insoluble at lower temperatures. In fact, constructs with truncated N-termini did not express at 18°C at all, and even the wild-type protein forms only inclusion bodies under such experimental settings (Fig. S6A-B). These data further stress the crucial role of the PfGSK3 N-terminus as well as its phosphorylation sites for folding and function of PfGSK3. As the N-terminally truncated PfGSK3 constructs are devoid of any phosphorylation, we propose that the N-terminal PfGSK3 domain, unique for *Plasmodium* species, is essential for PfGSK3 stability.

**Fig. 6.**
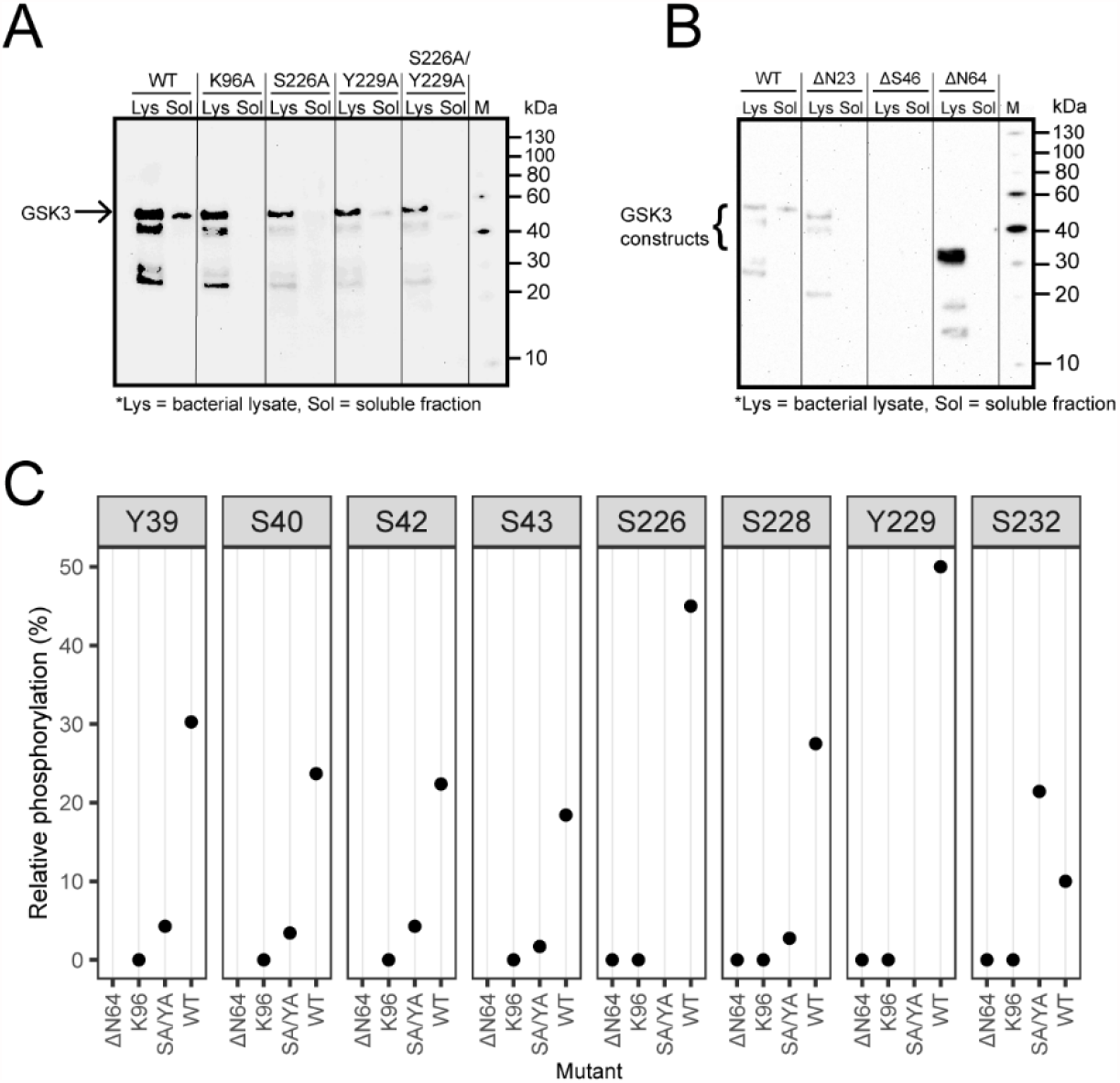
The N-terminus and phosphorylation is essential for PfGSK3. (A) Small-scale expression test of PfGSK3 mutants with inactivating mutations in the ATP binding site (K96A) and activation loop (Y226A and S229A). Whereas all protein constructs are expressed, only the wild type PfGSK3 is soluble, which indicates that the autophosphorylation of PfGSK3 is important for its solubility. (B) Small-scale expression test of PfGSK3 N-truncated constructs that start with residues N23, S46 or N64. In spite of a very strong expression of PfGSK3-N64, none of the proteins is soluble, indicating that the PfGSK3 N-terminus is crucial for the autophosphorylation process. The bacterial lysates (Lys) and their soluble fractions (Sol) were analyzed by Western blot with anti-His antibodies. (C) Analysis of phosphorylation of PfGSK3 mutants shows that the phosphorylation is completely lost (K96A, ΔN64) or reduced (S226A/Y229A) compared to the wild-type protein (WT).

### N-terminal phosphorylation fine-tunes the activity of PfGSK3

Provided that the PfGSK3 N-terminus is indispensable for its function, we suspected that this region could also be a potential site of PfGSK3 regulation, as it is in human GSK3β. Therefore, we analyzed the individual separated fractions from IEX chromatography (Fig. 1D) by LC/MS-MS. Here, we observed that the extent of phosphorylation of residues in the activation loop of PfGSK3 is comparable between the four fractions, but large differences in the N-terminal phosphorylation patterns are obvious (Fig. 7A). Specifically, all N-terminal residues are significantly less phosphorylated in the first fraction (F1), while in the second fraction (F2), the extend of phosphorylation of residues Y39 and S40 is strongly increased and the third and fourth fractions (F3 and F4) have generally higher level of N-terminal phosphorylation, which is most pronounced on residues Y39, S40, S42 and S43. Of note, only residue S40 is well conserved across *Plasmodium* species, but all other *Plasmodium* species contain multiple non-conserved N-terminal residues that are potential phosphorylation sites (Fig. S7A, F). To assess the relationship between the activity of PfGSK3 and its phosphorylation pattern, we made use of the differences in N-terminal phosphorylation between the individual IEX fractions and measured their kinase activity separately (Fig. 7B). PfGSK3 activity rises with an increase in N-terminal phosphorylation levels (from F1 to F4), supporting that phosphorylation is crucial for activity. Phosphorylation levels of residues S40, S42 and S43 significantly correlate with the PfGSK3 activity (Pearson R=0.6, p<0.05 in all cases, Fig. 7C-E), while this correlation is absent or insignificant for the other residues (Fig. S7B-F). These data support the hypothesis that phosphorylation of N-terminal PfGSK3 residues enhances PfGSK3 activity.

**Fig. 7.**
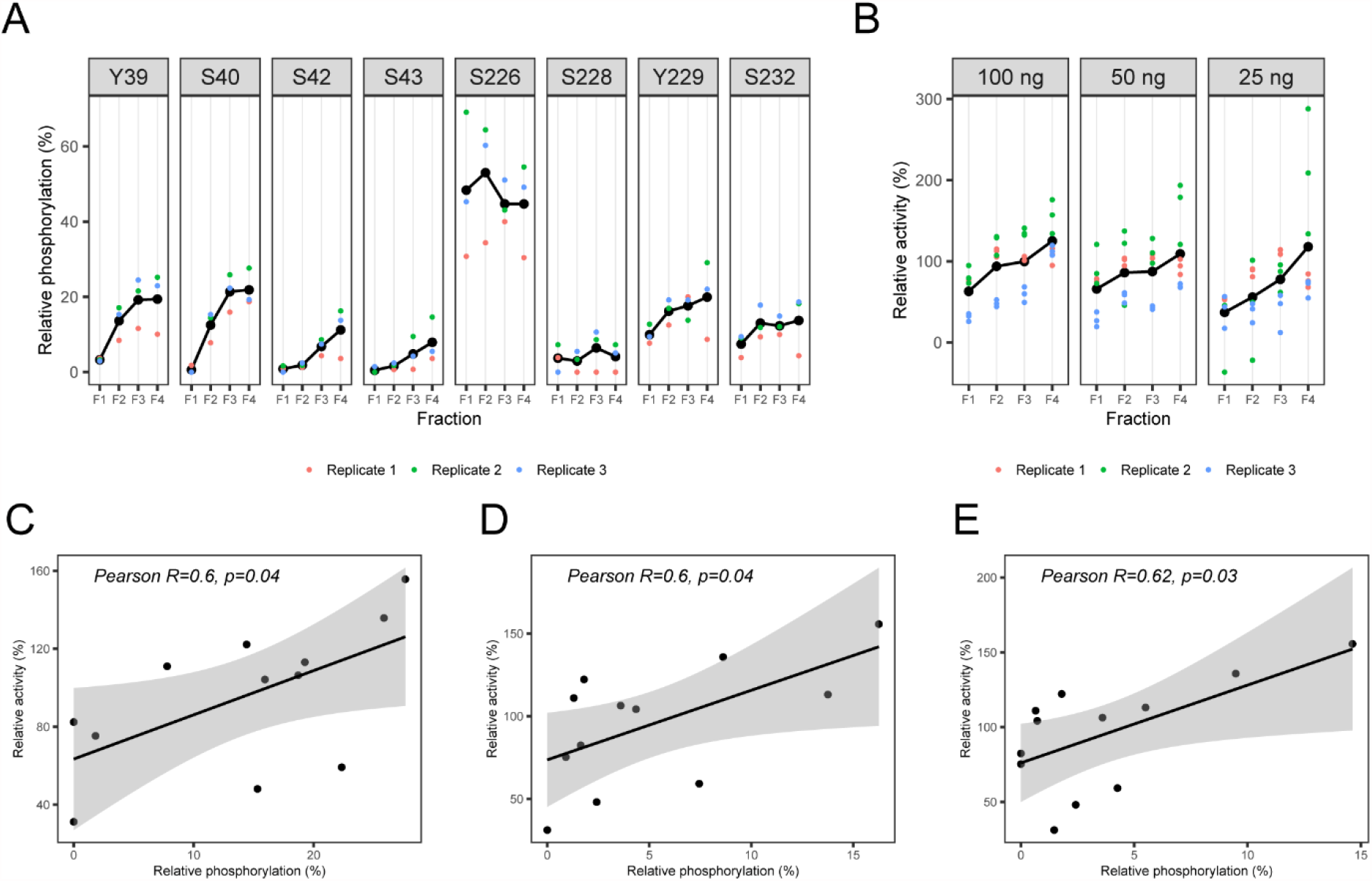
N-terminal phosphorylation promotes PfGSK3 activity. (A) Relative phosphorylation of selected residues in the individual PfGSK3 fractions separated by ion exchange chromatography. The relative phosphorylation represents the fraction of all identified residues that were also identified as phosphorylated by LC-MS/MS with Mascot score > 32 and MD score ≥ 5. The data show that the residues in the N-terminal domain are gradually more phosphorylated with increasing IEX elution volume, whereas the phosphorylation at the activation loop remains constant. The different colors represent three biological triplicates. (B) Activity of individual PfGSK3 fractions separated by ion exchange chromatography measured with different PfGSK3 amount (100, 50 and 25 ng). The data show a general trend towards higher activity with increasing phosphorylation in the N-terminal domain. The activity was measured in biological triplicates and at different PfGSK3 concentrations. (C-E) Correlation between the relative phosphorylation of the residues S40, S42 and S43 measured by mass spectrometry, respectively, and the relative activity of the same samples (triplicates of fractions F1-F4). The data were fitted with a linear model (black line), the gray background shows the 95% confidence interval.

### N-terminal phosphorylation affects the PfGSK3 structure

The phosphorylated residues at the PfGSK3 N-terminus are, based on structural homologs (24, 25), expected to be distant from the active site of the protein and it is therefore unclear how N-terminal phosphorylation regulates the protein activity. In the human counterpart, the phosphorylation of S9 prompts the binding of the N-terminus to a pocket for a priming phosphorylation in the substrate-binding region and reduces the binding affinity for substrates (24, 25). Hence, we investigated structural consequences of N-terminal phosphorylation in PfGSK3. Circular dichroism data on the least (F1) and most (F4) phosphorylated fractions show no pronounced differences in their secondary structure content (Fig. 8A). Thus, to investigate the impact of the N-terminal phosphorylation on the tertiary structure of PfGSK3, we used the *de novo* structure prediction server Robetta and the TrRefineRosetta method (70) to generate initial PfGSK3 models that were further fitted against the SAXS data measured on separated PfGSK3 IEX fractions F1 and F4 (Table 1, see Fig. 1D). Using the wild type PfGSK3 sequence as well as PfGSK3 sequences with phosphomimetic mutations (see Methods), we generated 25 models (Fig. 8A) that display similar structure of the core domains but are structurally diverse in both termini (Fig. 8A). We identified a subset of models that provide the best fits to the experimental SAXS data (Table 1, Fig. S8B) of fractions F1 or F4 (Fig. 8D). Overall, the models with a larger radius of gyration (see the point sizes in Fig. 8D) display lower χ^2^ (goodness-of-fit) values. Further refinement of these best-fitting extended models was conducted in the program CORAL (71) that enables assigning additional flexibility to existing models (Fig. 8E,H). This approach led to poorer fits when only C-terminus (residues 403-440) or both N-terminus and C-terminus (residues 1-64 and 403-440) were assigned flexibility (Fig. 8E,H). However, conducting refinement with a disordered C-terminus (residues 403-440) and a flexible stretch of residues between the N-terminus and the PfGSK3 core (residues 47-63, conformation B, Fig. 8E,H) led to two refined models displaying lower χ ^2^ values compared to the SAXS data from fraction F4. Ultimately, for both fractions, the models with the lowest χ^2^ values display α-helical structure at the N-terminus. This α-helix is extended in the less phosphorylated F1 fraction (Fig. 8B) and folds back in the more phosphorylated F4 fraction (Fig. 8C). The raw SAXS data support such conformational differences, as both the calculated radius of gyration *R*_*g*_ and maximum intra-particle distance *D*_*max*_ decrease from F1 to F4 fraction (*R*_*g*_ 3.4 vs 3.2 nm; *D*_*max*_ 13 vs 12 nm, respectively, see Table 1). The dimensionless Kratky plot data representation also suggests a more extended conformation for the F1 fraction (Fig. S8C), as the F1 curve broadens and shifts to higher *sR*_*g*_ values relative to F4. The refined models fit the SAXS data well (χ^2^ values of 1.19 and 1.06, respectively, Fig. 8F,G), but should be viewed cautiously as an average representation of the conformations present in solution that contains a mixture of differently phosphorylated species. In conclusion, the combinatorial approach indeed shows that upon phosphorylation, PfGSK3 becomes more flexible, enabling a conformational change in the N-terminus. This change presumably mediates the increased PfGSK3 activity upon phosphorylation.

**Table 1.** SAXS sample details, data acquisition parameters, structural parameters and atomistic modelling.

| Sample details |  |  |
| --- | --- | --- |
| Sample | PfGSK3 – F1 | PfGSK3 – F4 |
| Organism | *P. falciparum* |  |
| Source | *E. coli* (DE3) C41 |  |
| UniProt ID | O77344 |  |
| Extinction coefficient $\epsilon$ (at 280 nm, M<sup>-1</sup> cm<sup>-1</sup>) | 48250 | |
| Molecular weight from chemical composition (Da) | 52045.2 |  |
| Concentration (analysis or injection, mg/ml) | 10 |  |
| Solvent composition | 20mM Tris pH 8.0, 100mM NaCl, 0.5mM TCEP |  |
| SAS data collection parameters |  |  |
| Beamline | P12, DESY/EMBL, Hamburg (Germany) |  |
| Detector | Pilatus 6M |  |
| Energy (keV) | 10.0 |  |
| Sample-to-detector distance (mm) | 3000 |  |
| *q*-measurement range (Å<sup>-10</sup>) | 0.003-0.732 |  |
| Absolute scaling method | Relative to the scattering of pure water |  |
| Method for monitoring radiation damage | Frame comparison |  |
| Exposed time for frame | 900 s (900 x 1 s) | 900 s (900 x 1 s) |
| Mode | SEC-SAXS | SEC-SAXS |
| Sample temperature (°C) | 20 | 20 |
| Structural parameters |  |  |
| Guinier analysis |  |  |
| I(0) (cm<sup>-1</sup>) | 0.037±0.001 | 0.021±0.001 |
| R<sub>g</sub> (Å) | 34.3±0.4 | 31.5±0.2 |
| q-range (Å<sup>-1</sup>) | 0.01-0.04 | 0.01-0.04 |
| Fidelity (Quality of fit parameter, *AutoRg*) | 0.58 | 0.98 |
| Molecular weight (kDa)* |  |  |
| From V<sub>c</sub> | 66.2 | 48.7 |
| From MoW | 86.0 | 34.1 |
| P(r) analysis |  |  |
| I(0) (cm<sup>-1</sup>) | 0.037±0.001 | 0.021±0.001 |
| R<sub>g</sub> (Å) | 35.8±0.01 | 32.5±0.01 |
| D<sub>max</sub> (Å) | 130±10 | 115±10 |
| q-range (Å<sup>-1</sup>) | 0.01-0.50 | 0.01-0.25 |
| Total quality estimate | 0.62 | 0.82 |
| Porod volume x 10<sup>3</sup> (Å<sup>3</sup>) | 105 | 101 |
| **Atomistic modelling\*\*** | CRY SOL with constant subtraction and maximum order of harmonics equal to 50 | CORAL hybrid rigid body modeling |
| Structures | Robetta model WLT (4) |  |
| q-range for modelling (Å<sup>-1</sup>) | 0.01-0.50 | 0.01-0.58 |
| χ<sup>2</sup> | 1.19 | 1.06 |
| Predicted R<sub>g</sub> (Å) | 35.3 |  |
| Starting crystal structures |  | Robetta model 3SE (4) |
| Flexible residues |  | 47-63, 403-440 |
| **SASBDB IDs for data and models** |  |  |
|  | SASDL77 | SASDL87 |

\* calculated using the appropriate functions in ATSAS (91).
\*\* The values for the best model for each respective dataset are shown.

**Fig. 8.**
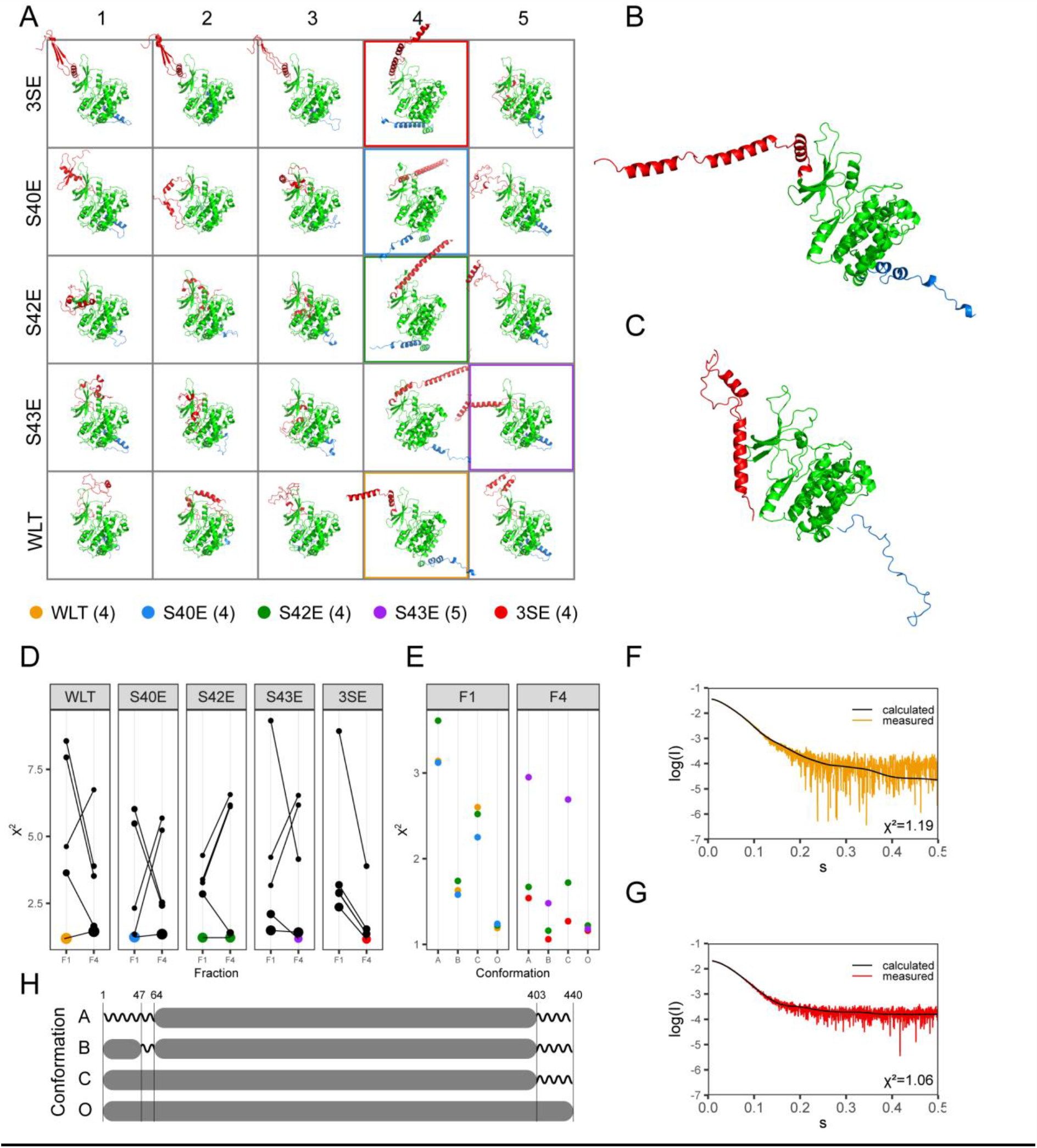
The N-terminus of PfGSK3 changes the structure upon phosphorylation. (A) Structural models of PfGSK3 predicted by the Robetta server. Five different sequences were used as an input for the prediction and five models were predicted for each sequence. N-termini (residues 1-65) are red, the core domains green and C-termini (residues 403-440) blue. (B) The Robetta model that fits the SAXS data from F1 fraction best display an extended N-terminal helix. (C) The Robetta model refined against SAXS data from fraction F4 using CORAL with the best fit to the data. The N-terminal helix of this model folds back towards the core of PfGSK3. (D) *Χ*^*2*^ values of PfGSK3 structures modelled based on five different sequence modifications (WLT, S40E, S42E, S43E and 3SE) compared to SAXS data recorded for fractions F1 or F4. The identical models are connected with lines. The size of each points correlates with the radius of gyration of the particular model. Three models with lowest *Χ*^*2*^ values for each F1 and F4 are colored (see legend). (E) *Χ*^*2*^ values of the chosen models refined with CORAL using different strategies, enabling flexibility at either the N-terminus (residues 1-63, strategy A), C-terminus (residues 403-440, strategy C), or at both the C-terminus and residues 47-63 (strategy B). *Χ*^*2*^ values of the original models are shown under “O”. (F) The SAXS data measured on the PfGSK3 fraction F1 with the fit of theoretical scattering calculated from the best fitting model (*Χ*^*2*^=1.19). (G) The SAXS data measured on the PfGSK3 fraction F1 with the fit of theoretical scattering calculated from the best fitting CORAL-refined model (*Χ*^*2*^=1.06). (H) Graphical summary of the strategies used for model refinement against SAXS data using CORAL. The gray box symbolizes that the structure of the model was maintained and the zigzag line symbolizes that the structure was replaced with disordered residues by CORAL.

## Discussion

Identification and characterization of novel drug targets in malaria remain an important goal as drug-resistant parasites are a persisting threat in the fight against malaria. PfGSK3 has been identified as a potential drug target since its inhibitors also possess potent and selective anti-plasmodial activity. Understanding its function is therefore important for further improvement of these inhibitors.

Here, we have shown that PfGSK3 is expressed in *E. coli* as a folded and functional protein with a heterogeneous phosphorylation pattern. We assume that the heterogeneity originating from this diverse phosphorylation is the main reason why we were not able to grow PfGSK3 crystals for detailed structural characterization (72). Moreover, the PfGSK3 C-terminus, predicted to be disordered, and its unusual N-terminus represent additional challenges in protein crystallization. As the expression of inactive PfGSK3 protein (K96A, S226A, Y229A) as well as the removal of heavily phosphorylated N-terminus result in insoluble protein that lacks phosphorylation, other protein constructs and strategies will likely need to be considered for the structure determination of PfGSK3 in the future. At the same time, these results highlight the importance of both PfGSK3 phosphorylation and the necessity of an intact N-terminus for protein stability.

Our mass spectrometry analysis of the phosphorylation pattern on recombinant PfGSK3 confirmed a number of PfGSK3 phosphorylation sites known from previous phosphoproteomics studies (S31, S62, S226, S228, Y229, S232) (37–39, 62–65). The additional phosphorylation sites at the N-terminus (Y39, S40, S42 and S43) might have occurred due to the heterologous expression, but could also hint towards additional phosphorylation sites present *in vivo* under certain conditions. In general, these phosphorylation sites are reminiscent of those identified in human GSK3β. In GSK3β, the autophosphorylation of Y216 in the activation loop is thought to maintain the constitutive activity of the protein (27, 29), while phosphorylation of N-terminal S9 by other kinases, such as PKA and PKB, inhibits the enzyme (6, 30). Although the role of the activation loop autophosphorylation is likely conserved between the human and *P. falciparum* proteins, the role of N-terminal phosphorylation seems to be different.

First, the N-terminal sequence is not conserved between the two homologous proteins, with *Plasmodium* species displaying an unusual extension with a number of potential phosphorylation sites. Second, whereas N-terminal phosphorylation of S9 in human GSK3 is mediated by other kinases, the phosphorylation of N-terminal residues in PfGSK3 is the result of autophosphorylation, as we show here. Last, while phosphorylation of S9 in human GSK3 inhibits its activity, our work illustrates that the increasing amount of phosphorylation on the PfGSK3 N-terminus results in an increased enzymatic activity.

Our structural investigation using the combination of structure prediction and its validation using experimental SAXS data provides further insights into how the N-terminal phosphorylation affects PfGSK3 structure. Two SAXS datasets originating from two differently phosphorylated IEX fractions of PfGSK3 directed the modelling towards PfGSK3 conformations with an α-helical N-terminus in both cases. However, this helix extends away from the active side of PfGSK3 in the less phosphorylated fraction, or folds back towards the center of the protein in the more phosphorylated fraction. It is possible that the “folded back” conformation (Fig. 8C) could be mediated by an N-terminal intramolecular coiled-coil interaction induced by phosphorylation, as the probability of coiled-coil formation increases with the increasing number of N-terminal PfGSK3 phosphomimetic mutations *in silico* (Fig. S8D). Interestingly, human GSK3β undergoes a similar conformational change, where upon phosphorylation of S9, its N-terminus binds in its substrate-binding region and competitively inhibits its activity (24, 25). However, N-terminal phosphorylation of PfGSK3 leads to an increase in enzymatic activity, suggesting a different regulatory mechanism. The conformational change upon phosphorylation might position the very N-terminus of PfGSK3 close to its core domains, potentially enabling intramolecular interactions that could enhance its activity. Alternatively, the N-terminal extension might interact with the substrates to guide them to the PfGSK3 active site. In both cases, the conformational change would be driven by phosphorylation of residues 39-43, whereas the interacting residues are possibly confined to the very N-terminus (residues 1-20) that is conserved in *Plasmodium* species (Fig. S6A,F). It should be noted that the N-termini of PfGSK3 homologues are not well conserved even among the representatives of *Apicomplexa*; however, a number of residues that can be phosphorylated could still enable a similar mode of regulation as we have described here (Fig. S7A,F). Indeed, S33 of *Plasmodium berghei* GSK3 (PBANKA_0410400), equivalent to S40 in PfGSK3, was found to be phosphorylated in early gametogenesis (73). Residues equivalent to S40 of PfGSK3 are conserved across all *Plasmodium* species and residues that can be phosphorylated are also present in the vicinity of S40 in most *Plasmodium species* (equivalent to residues 38, 39 and 42 in *P. falciparum*), suggesting that the mechanism of regulation *via* N-terminal phosphorylation is preserved in all *Plasmodium* GSK3 proteins.

We have shown that this N-terminal phosphorylation, as well as the phosphorylation of the activation loop, are the result of PfGSK3 exhibiting autophosphorylation. This result confirms data in a recent publication, where the authors suspected that a high background in their phosphorylation assay could be attributed to autophosphorylation (74). Autophosphorylation of human GSK3β occurs directly after protein synthesis while the protein is associated with chaperones, such as HSP90 (27). Interestingly, a major contaminant during our PfGSK3 purification process was *E. coli* DnaK that shares 48-57% identity with various *P. falciparum* Hsp70 proteins, indicating that chaperone-assisted autophosphorylation might be conserved across these taxa. Moreover, PfGSK3 was less soluble and had lower expression levels at lower temperatures (25°C and 18°C), at which the chaperone levels in *E. coli* are also reduced (75). This further supports that chaperons could be crucial interaction partner for the first stages of PfGSK3 autophosphorylation, which are likely crucial for its solubility.

Additionally, we have revealed another means of PfGSK3 regulation: the divalent heavy metal ions, such as zinc and copper, inhibit the activity of PfGSK3 by inducing the formation of heterogeneous high-MW particles. The ionic radii of these cations are in a small range of 1.09-1.21 Å, indicating that the effect is mediated through a specific binding site. The fact that the protein becomes more sensitive to the metal ions only after the cleavage of the N-terminal affinity tag indicates that this potential binding site could be located in close proximity to the N-terminus of PfGSK3. Indeed, a partially overlapping set of divalent cations inhibiting human GSK3β can be found in the literature, including lithium, beryllium, zinc, copper and mercury (17, 76, 77). Lithium and beryllium ions inhibit GSK3 by competing with magnesium ions that mediate ATP binding (77–79), but to our knowledge, they have not been shown to induce the formation of high-MW particles. To this date, there has been no structure of a GSK3 homolog with a bound heavy metal ion deposited in the PDB database, and thus the inhibition mechanism of heavy metal cations remains unknown. Importantly, as the high-MW particles can dissociate back into active PfGSK3 monomers, it cannot be excluded that this type of regulation is applied *in vivo*. The concentration of zinc in *Plasmodium* cytosol peaks at the late blood stages and should be sufficient to induce the formation of high-MW PfGSK3 species (80). Moreover, the concept of “auto-inhibitory polymerization” has already been described in detail for human CK2 (81), further underlining the possible similarities between PfCK2 and PfGSK3. It remains to be evaluated experimentally *in vivo* whether the PfGSK3 high-MW particles play a role in its native environment.

In conclusion, our work provides new insights into the function of PfGSK3 and its regulation by N-terminal phosphorylation and heavy metals, highlighting that phosphorylation influences the structure of the PfGSK3 N-terminus that in turn results in increased kinase activity.

## Experimental procedures

### Cloning and mutagenesis

The vector with N-terminally His-tagged PfGSK3 was generated by PCR amplification of the GSK3 coding sequence from *P. falciparum* cDNA followed by Ligation Independent Cloning into HindIII/KpnI-cleaved plasmid pOPIN F (82) using the In-Fusion HD EcoDry Cloning Kit (Takara Clontech) according to the manufacturer’s instructions. The mutants S226A, Y229A and S226A/Y229A were generated by overlap extension PCR amplification from the original vector and Ligation Independent Cloning as described above. The wild-type protein and the mutant K96A cloned in pET28a vector were ordered from GenScript. The N-terminally truncated constructs were cloned by amplifying the sequence from the original vector and subcloning into BsaI-cleaved plasmid pNIC28_Bsa4 by SLiCE cloning (83). The construct pSLI-GSK3-GFP-glms was generated by PCR-amplification of 1013 bp of the C-terminal GSK3 coding sequence from genomic DNA using primers GSK3_NotI_sense_SLI and GSK3_AvrII_as_fored_int and cloning into pSLI-GFP-glms (84, 85) using NotI/AvrII restriction sites.

### PfGSK3 expression and purification

*E. coli* C41(DE3) culture transformed with the PfGSK3 coding vector was grown in TB medium supplemented with 0.1 mg/ml ampicillin at 37°C, induced at OD_600nm_=0.7 with 0.5 mM IPTG and harvested after 4 hours. The pellets were resuspended in 5 mL of lysis buffer (20mM NaP pH 7.5, 300mM NaCl, 15mM imidazole, 5% glycerol, 0.5mM TCEP, 1 mg/ml lysozyme, 5 U/ml DNase, 1 Roche protease inhibitor tablet/100mL) per 1 g of wet weight culture. The suspension was passed three times through an emulsifier at 15 000 psi, centrifuged at 19 000 x g and the supernatant was loaded on a pre-equilibrated His-Trap column. The column was washed with lysis buffer supplemented with 40 mM imidazole and the protein was eluted at increasing imidazole concentrations. The protein containing fractions were concentrated (10 kDa cut-off concentrator) with the addition of 1 mg of 3C protease per 3 L of culture volume and further separated on a Superdex 200 HiLoad column (GE Healthcare) in 50mM Tris pH 8.0, 20mM NaCl, 0.5mM TCEP. The protein containing fractions were concentrated again, loaded on ResourceQ anion exchange column (GE Healthcare) and eluted with increasing concentration of NaCl. Fractions representing different PfGSK3 species were pooled together, concentrated, dialyzed against the analysis buffer (typically 20mM Tris pH 8.0, 100mM NaCl, 0.5mM TCEP) and flash-frozen in liquid nitrogen and stored at −80°C. All purification steps were performed at 4°C.

### Expression and purification of insoluble PfGSK3 constructs

The insoluble PfGSK3 constructs (K96A, S226A/Y229A, ΔN64) were expressed as described for the wild-type PfGSK3. The pellets of harvested bacteria were resuspended in 5 mL of lysis buffer (20mM NaP pH 7.5, 300mM NaCl, 15mM imidazole, 5% glycerol, 0.5mM TCEP, 1 mg/ml lysozyme, 5 U/ml DNase, 1 Roche protease inhibitor tablet/100mL) per 1 g of wet weight culture. The suspension was passed three times through an emulsifier at 15 000 psi, centrifuged at 19 000 x g and the pellet was dissolved in a denaturating buffer (20mM NaP (pH 7.5), 300mM NaCl, 25mM imidazole, 5% glycerol, 6M urea, 0.5mM TCEP) and incubated with 1 mL NiNTA bead slurry (Sigma) for 1h, 4°C. The beads were subsequently washed twice with 10 ml of the same buffer and subsequently eluted with 5 ml of the same buffer containing extra 250 mM imidazole. The protein was concentrated (10 kDa c/o) and the sample for mass spectrometry was separated by SDS-PAGE gel.

### Small-scale expression tests

4 mL of TB supplemented with ampicillin was inoculated wit pre-cultured *E. coli* C41(DE3) that had been transformed with the tested plasmids. The cultures were grown at 37°C until the OD_600nm_ =0.6 and further either induced with 0.5mM IPTG and grown for 4 hours at 37°C or 25°C, or induced with 0.1mM IPTG and grown for 16 h at 18°C. OD_600nm_ was measured and culture volume corresponding to OD_600nm_ of 2.0/mL was harvested. The pellets were resuspended in 400 µl of lysis buffer (lysate fractions), then lysed by 10 min of vortexing with 100 μl glass beads and the cell debris were removed by centrifugation (soluble fractions). 5 µl of each fraction was mixed with 10 µl of loading dye and run on SDS-PAGE in duplicates. One gel of each set of samples was blotted on PVDF membrane (*BioRad*) The membranes were blocked by 1% BSA in TBS-T, washed, incubated for 1h with a HisProbe-HRP conjugated antibody (ThermoFisher), washed with TBS-T and the blot was developed using Super Signal West Pico Substrate (ThermoFisher) and Super Signal West Femto Substrate (ThermoFisher) in a 1:10 ratio.

### Glutaraldehyde cross-linking

1 mg/ml PfGSK3 in PBS and 0.5mM TCEP was mixed with 8x molar excess of either EDTA or ZnCl_2_ and 0.01% glutaraldehyde. The reaction was quenched with 330mM Tris (final concentration) and SDS-PAGE loading dye and SDS-PAGE was run for 150 minutes at 150V. The gels were stained with Direct Blue stain.

### Thermal shift assay

The stability of PfGSK3 under different buffer conditions and additives was measured by nanoDSF (Prometheus NT.48, NanoTemper Technologies, GmbH). The protein concentration was adjusted to 1 mg/ml. 10 μl of samples were loaded in the glass capillaries and heated from 20°C to 90°C with a heating rate of 1°C/min. The fluorescence signals with excitation wavelength of 280 nm and emission wavelengths of 330 and 350 nm were recorded and the melting temperature was calculated as maximum of the derivative of the fluorescence at 330 nm.

### Circular dichroism

Circular dichroism was measured on a Chirascan CD spectrometer (Applied Photophysics). The protein concentration was adjusted to 1 μM by 2 mM NaP (pH 7.5), 4 mM NaCl, 0.05 mM TCEP prior to the measurement. The circular dichroism spectrum was measured 10x between 185 nm and 260 nm with 1 nm step in 1 mm quartz cuvette and analyzed by the CDSSTR algorithm (86, 87) using DichroWeb (88).

### Analytical size exclusion chromatography

The analytical size exclusion chromatography was performed on an Agilent Bio-LC system using a home-packed Superose 5/150 column (25 μl sample) or a Superose 6 Increase 10/300 column (100 μl sample). PBS supplemented with 0.5mM TCEP was used as a mobile phase and the resulting chromatograms were recorded using the as fluorescence signal with excitation wavelength of 280 nm and emission wavelength of 350 nm as readout.

### Autophosphorylation IEX assay

0.5 mg/ml of PfGSK3 was incubated at 37°C with 2mM ATP or cAMP and 10mM MgCl_2_ overnight. The samples were subsequently dialyzed at 4°C overnight in 50mM Tris pH 8.0, 20mM NaCl, 0.5mM TCEP. The samples were analyzed by analytical ion exchange chromatography using a Resource Q column (GE Healthcare) and the 1260 Infinity Bio-inert high-performance liquid chromatography system (Agilent Technologies) at 10 °C. The system and column were equilibrated in 50mM Tris pH 8.0, 20mM NaCl, 0.5mM TCEP. 100 µl of sample was injected and eluted with an increasing concentration of NaCl. The system was run at 0.2 ml/min ad the elution profile was analyzed by UV fluorescence detector with absorbance at 280 nm and emission at 350 nm.

### Small angle X-ray scattering

The SAXS data were collected at the P12 BioSAXS Beamline at the PETRA III storage ring (c/o DESY, Hamburg, Germany) (89). The concentrated samples were dialyzed against (20 mM Tris pH 8.0, 100 mM NaCl, 0.5 mM TCEP overnight at 4°C and the concentration was adjusted to 10 mg/ml The X-ray scattering data were measured in an on-line SEC-SAXS mode, using a SD200 Increase 5/150 column (GE Healthcare) at 0.4 ml/min with 1 frame recorded per second (Table 1). The data were automatically processed using the SASFLOW pipeline (90) and further analyzed using the ATSAS suite (91) of programs to determine the molar mass, radius of gyration and real-space distance distribution. Scattering curves were computed from models using CRYSOL (92) with a constant subtraction and maximum order of harmonics set to 50. The rigid body modelling performed using CORAL (71), setting different parts of models yielding best CRYSOL fits as disordered. These were residues 1-65 and 403-440 in conformation A, resides 47-65 and 403-440 in conformation B and residues 403-440 in conformation C. The raw data were uploaded to SASBDB under accession numbers SASDL77 and SASDL87.

### Kinase assay

To investigate enzymatic activity of recombinant PfGSK3 a commercial luminescence-based kinase assay (KinaseGlo Plus, Promega) was used as previously described (93). Briefly, 20 ng PfGSK3, 12 µM GS-1 peptide substrate (residues 636-661 of the human glycogen synthase 1; sequence YRRAAVPPSPSLSRHSSPHQ(pS)EDEEE; pS = prephosphorylated serine, Promega) and 6 µM ATP (UltraPure, Promega) in the kinase reaction buffer (40 mM Tris-HCl pH 7.5; 20 mM MgCl_2_; 0.1 mg/mL BSA) were used as standard reaction mix (final reaction volume 5 µl). The kinase reaction was performed for 30 min at 30°C followed by incubation with KinaseGlo reagent according to the manufacturer’s instructions. The reaction was transferred to a solid white 384-well plate (NUNC, ThermoFisher) and the luminescence signal was measured in an EnVision Multilabel Plate Reader (PerkinElmer, Integration time 0.5 sec/well). To investigate reaction kinetics, kinase and substrate concentrations were varied between 0 and 100 ng or 0 and 15 µM respectively. To investigate autophosphorylation, reactions were performed with kinase concentrations between 0 and 500 ng, 30 µM ATP and without a substrate.

### *P. falciparum* culture

Parasites were cultured in 5% 0^+^ human erythrocytes (Blood bank, Universitätklinikum Hamburg Eppendorf) in RPMI medium supplemented with 0.5% Albumax at 37°C in an atmosphere of 1% O_2_, 5% CO_2_ and 94% N_2_ as described previously (94). Parasitemia was monitored by Giemsa-staining of methanol-fixed blood smears. The parasite strain 3D7 was used in this study (95).

### Parasite imaging

For live cell imaging parasite cultures were pelleted by centrifugation at 500x g for 1 min. Sedimented cells were resuspended in an equal volume of the supernatant, 5 µl of the sample were placed on a microscopy slide and covered with a cover slip. Fluorescence microscopy images were taken with a Leica DM6 B microscope equipped with Leica DFC9000 GT camera using an 100x immersion oil objective. Images were aquired with Leica Application Suite X (LAS X) software and processed using Adobe Photoshop CS2. Nuclei were stained with 1 µg/ml DAPI (Biomol) in RPMI at 37°C for 15 min.

### Generation of transgenic parasite lines

For Transfection, mature schizonts were enriched using 60% percoll and electroporated with 50 µg of plasmid DNA using a Lonza Nucleofector II (96). Transfectants were seleceted with 4nM WR99210 (Jacobus Pharmaceuticals). Stable integrant cell lines were generated as previously described (84). Briefly, parasites carrying the WR99210-selected episomal plasmid were cultivated in the presence of 400 µg/ml G418 (Sigma) to select for integrants with the desired modification of the genomic locus. Correct genomic integration of the construct was confirmed by PCR using genomic DNA from G418-selected integrants and 3D7 wildtype. Genomic DNA was isolated using QIAamp DNA Mini Kit. For PCR analysis primers specific for 3’ and 5’ integration as well as wildtype locus of PfGSK3 were used.

### Purification of GSK3-GFP from *P. falciparum*

For purification of GSK3-GFP, parasites were synchronized by treatment with 5% sorbitol for 10 min at 37°C and subsequently cultured in a volume of 50 ml with 5% hematocrit. Culture medium was changed daily until parasitemia had reached 8-10%. Erythrocytes were lysed with 0.03% saponin, isolated parasites were washed three times with ice cold PBS and resuspended in 2 ml lysis buffer (50 mM Tris-HCl pH 7.5, 150 mM NaCl, 0.5 mM TCEP, 1x protease inhibitor mix, 50 U/mL DNase). The sample was triturated five times through a 27G-needle (Braun) and further lysed by three cycles of freezing at −80°C followed by thawing on ice. The lysates were supplemented with 1% DDM and 0.1% CHS and incubated for 3 hours at 4°C. Subsequently, the samples were mixed with 30 µl of GFP-Trap bead slurry (ChromoTek) and incubated on a rotatory wheel for 3 hours at 4°C. The beads were then washed with 1 ml of 150 mM NaCl, 10 mM Tris, protease inhibitor mix (Roche), 0.1% DDM, 0.01% CHS pH 7.5 and with 1 ml of 300 mM NaCl, 10 mM Tris, protease inhibitor mix, 0.02% DDM, 0.002% CHS pH 7.5. The trapped proteins were eluted with 30 µl of 6% SDS, 250 mM Tris-HCl, pH 6.8, 71 mM TCEP at 95°C.

### Sample preparation for mass spectrometry

In-solution samples from parasite cell cultures were processed as previously described (97). To assess the phosphorylation status of the recombinant PfGSK3 (purified protein, separated IEX fractions, insoluble constructs), each sample was separated on SDS-PAGE gel. Coomassie-stained bands were excised, chopped into small pieces and transferred to 0.5 ml Eppendorf tubes. For all following steps, buffers were exchanged by two consecutive 15 min incubation steps of the gel pieces with 200 µl of acetonitrile (ACN), whereby the ACN was removed after each step. Proteins were reduced by the addition of 200 µl of a 10 mM DTT solution in 100 mM ammonium bicarbonate (AmBiC, Sigma Aldrich, A6141) and incubation at 56°C for 30 min. Proteins were alkylated by the addition of 200 µl of 55 mM chloroacetamide (CAA)100 mM AmBiC solution and incubation for 20 min in the dark. A 0.1 µg/µl stock solution of trypsin (Promega, V511A) in trypsin resuspension buffer (Promega, V542A) was diluted with ice-cold 50 mM AmBiC buffer to achieve a final concentration of 1 ng/µl. 50 µl thereof were added to gel pieces, which were incubated first for 30 min on ice and then over night at 37°C. Gel pieces were sonicated for 15 min, spun down and the supernatant was transferred into a glass vial (VDS Optilab, 93908556). Remaining gel pieces were washed with 50 µl of an aqueous solution of 50% ACN and 1% formic acid and sonicated for 15 min. The combined supernatants were dried in a speedvac and reconstituted in 10 µl of an aqueous solution of 0.1% (v/v) formic acid.

### Mass spectrometry data acquisition

The mass spectra of the initial phosphorylation site assessment of PfGSK3 (Fig. SB) was performed as follows: An UltiMate 3000 RSLC nano LC system (Dionex) fitted with a trapping cartridge (Precolumn C18 PepMap100, 5 mm, 300 μm i.d., 5 μm, 100 Å) and an analytical column (Acclaim PepMap 100. 75 × 50 cm C18, 3 mm, 100 Å) was used. Trapping was carried out with a constant flow of trapping solution (0.05% trifluoroacetic acid in water) at 30 µL/min onto the trapping column for 6 minutes. Subsequently, peptides were eluted via the analytical column running solvent A (0.1% formic acid in water) with a constant flow of 0.3 µL/min, with increasing percentage of solvent B (0.1% formic acid in acetonitrile) from 2% to 4% in 4 min, from 4% to 8% in 2 min, then 8% to 28% for a further 37 min, in another 9 min. from 28%-40%, and finally 40%-80% for 3 min followed by re-equilibration back to 2% B in 5 min. The outlet of the analytical column was coupled directly to an Orbitrap QExactive™ plus Mass Spectrometer (Thermo) using the Nanospray Flex™ ion source in positive ion mode. The QExactive plus was operated in positive ion mode with a spray voltage of 2.2 kV and capillary temperature of 275°C. Full scan MS spectra with a mass range of 350–1400 m/z were acquired in profile mode using a resolution of 70,000 (maximum injections time of 100 ms, AGC target was set to 3×10^6^ ions). Precursors were isolated using the quadrupole with a window of 1.7 m/z and fragmentation was triggered by HCD in fixed collision energy mode with fixed collision energy of 28%. MS2 spectra were acquired with the Orbitrap with a resolution of 17.500 and a max injection time of 70 ms.

The mass spectra of all other experiments were recorded as follows: Peptides were analyzed by LC-MS/MS on an Orbitrap Fusion Lumos mass spectrometer (Thermo Scentific) as previously described (98). To this end, peptides were separated using an Ultimate 3000 nano RSLC system (Dionex) equipped with a trapping cartridge (Precolumn C18 PepMap100, 5 mm, 300 μm i.d., 5 μm, 100 Å) and an analytical column (Acclaim PepMap 100. 75 × 50 cm C18, 3 mm, 100 Å) connected to a nanospray-Flex ion source. For the detection of posttranslational modified peptides, peptides were loaded onto the trap column at 30 µl per min using solvent A (0.1% formic acid) and peptides were eluted using a gradient from 2 to 80% Solvent B (0.1% formic acid in acetonitrile) over 60 min at 0.3 µl per min (all solvents were of LC-MS grade). The Orbitrap Fusion Lumos was operated in positive ion mode with a spray voltage of 2.4 kV and capillary temperature of 275 °C. Full scan MS spectra with a mass range of 375–1200 m/z were acquired in profile mode using a resolution of 120,000 (maximum injections time of 50 ms, AGC target was set to Standard with a max injection time of 50 ms. Precursors were isolated using the quadrupole with a window of 1.2 m/z and fragmentation was triggered by HCD in fixed collision energy mode with fixed collision energy of 34%. MS2 spectra were acquired with the Orbitrap with a resolution of 15.000 and a max injection time of 54 ms.

### Mass spectrometry data analysis

Acquired data were analyzed using IsobarQuant (99) and Mascot V2.4 (Matrix Science) using a reverse UniProt FASTA database from *E. coli* (UP000000625) or *Plasmodium falciparum* (UP000001450) including common contaminants. The following modifications were taken into account: Carbamidomethyl (C, fixed), Acetyl (K, variable), Acetyl (Protein N-term, variable), Oxidation (M, variable), as well as the variable modification Phospho (ST) and Phospho (Y). The mass error tolerance for full scan MS spectra was set to 10 ppm and to 0.02 Da for MS/MS spectra. A maximum of 2 missed cleavages were allowed. A minimum of 2 unique peptides with a peptide length of at least seven amino acids and a false discovery rate below 0.01 were required on the peptide and protein level (100). Prior performing the search, the protein sequences corresponding to the expressed PfGSK3 constructs were added to the database for the expression host. The raw data have been uploaded to the PRIDE database with the dataset identifier PXD026387.

### Correlation analysis

Each biological replicate of mass spectrometry experiment was paired with the corresponding biological replicate of the activity measurement. The relative phosphorylation of a residue was calculated as a ratio of number of time the residue was identified as phosphorylated (Mascot score >32 and MD score >=5) to number of times the residue was identified in the biological sample. The relative activity was calculated as buffer-subtracted readout of the luciferase assay relative to control. The strength of the linear association was calculated as Pearson correlation coefficient using R (101).

### Negative stain electron microscopy

4 µl of 0.01 mg/ml protein solution was applied to carbon-coated Cu/Pd grids and incubated for 1 min. Excess liquid was removed with the filter paper (Whatman), and the grid was washed twice with water for 5 s and 30 s. 4 µl of 2% uranyl acetate were applied twice for 5 s and 30 s. The micrographs were collected on a Talos L120C (CSSB Cryo-EM multiuser facility) with 70 000x magnification.

### Bioinformatics

The homologous protein sequences were aligned with the program MAFFT (102) and depicted in Jalview. The secondary structure was predicted by Jpred (103). The PfGSK3 structures were predicted using the Robetta server and the TrRefineRosetta method (70). To simulate the different N-terminally phosphorylated species present in the protein fractions, we generated five models for each of the following PfGSK3 sequences: wild-type sequence, three sequences with a single phosphomimetic mutation (S40E, S42E, S43E) and one with a triple mutation S40E/S42E/S43E (abbreviated SE3). Per-residue local error estimates of the models calculated by Robetta (104) are written in the B-factor column of the PDB files. For phylogenetic analysis, protein sequences with highest identity scores were identified for individual organisms with the blastp suite of NCBI (105), the sequences were aligned with the ClustalW method and the R package msa (106), and the phylogenetic tree was calculated in PhyML with LG +G+I+F method (107). The phylogenetic tree with multiple sequence alignment was plotted using R package ggtree (108).

## Data availability

The mass spectrometry data are available in PRIDE database with dataset identifier PXD026387. The SAXS data have been deposited in the SASBDB, accession numbers SASDL77 and SASDL87. All other raw data are available on Figshare (doi: 10.6084/m9.figshare.14806887).

## Supporting information

This article contains supporting information.

## Acknowledgments

We thank the Sample Preparation and Characterization facility of EMBL Hamburg for support in nanoDSF, CD and DLS measurements. We also thank the Proteomics Core Facility of EMBL for the support in mass spectrometry sample preparation, measurements and analysis. We thank Maxime Killer and Vadim Kotov for the help with EM data acquisition and processing. Part of this work was performed at the CryoEM Facility at CSSB, supported by the UHH and DFG grant numbers (INST 152/772-1|152/774-1|152/775-1|152/776-1|152/777-1 FUGG). This work was supported by a grant from the Joachim Herz foundation (Nr. 800026) and the BMBF (grant number: 05K18YEA). AA is grateful for support by a Jürgen Manchot Stiftung fellowship.

## Abbreviations

AMA1: *Plasmodium falciparum* Apical membrane Antigen 1
GSK3β: Glycogen Synthase Kinase beta
IEX: ion exchange chromatography
IMAC: immobilized metal affinity chromatography
high-MW: high molecular weight
PfGSK3: *Plasmodium falciparum* Glycogen Synthase Kinase
SEC: size exclusion chromatography

