## Supplementary information for "N-terminal phosphorylation regulates the activity of Glycogen Synthase Kinase 3 from *Plasmodium falciparum*"

### Supporting information

Supplementary figure 1

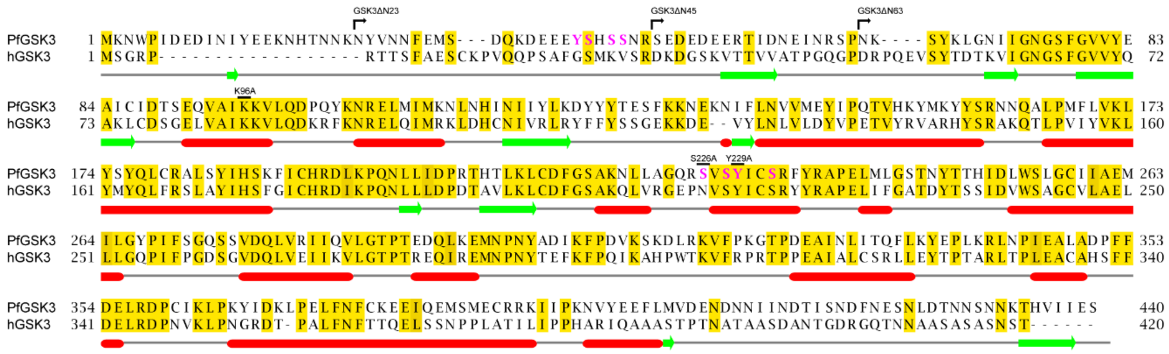


Sequence alignment of PfGSK3 with human GSK3. The conserved residues are highlighted in yellow. Residues mutated in this study and the N-truncated constructs are indicated. Residues that are phosphorylated are shown in magenta. The PfGSK3 secondary structure prediction by JPred (109) is indicated below the sequence. The sequences were aligned using MAFFT (102) and plotted in Jalview (103).

Supplementary figure 2

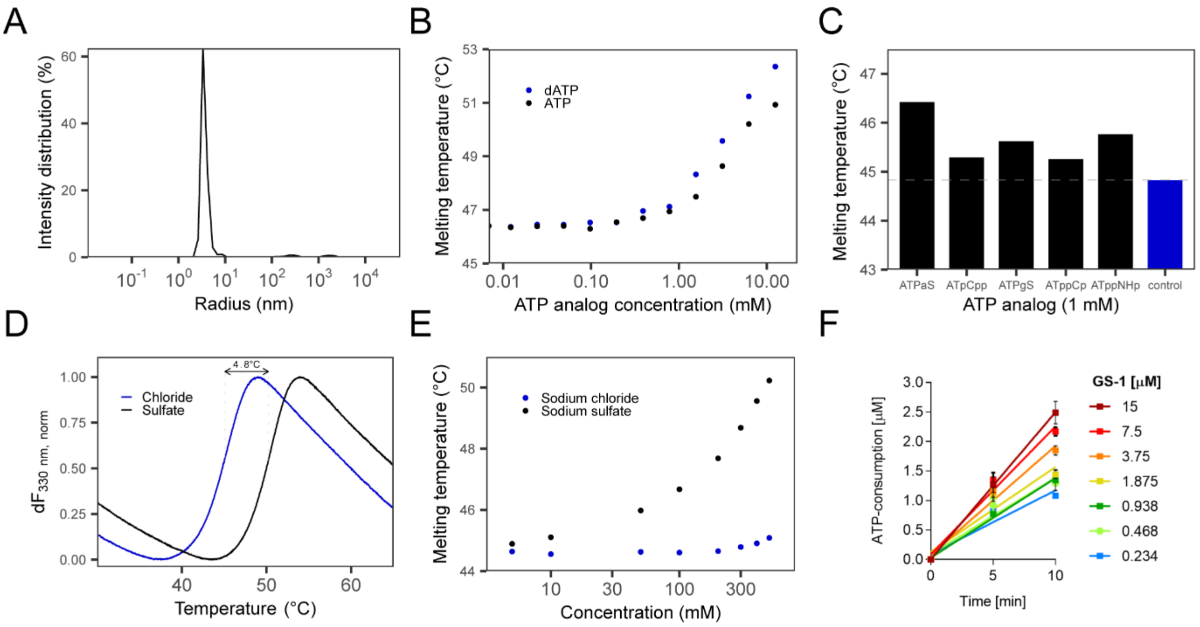


(A) Analysis of dynamic light scattering data of PfGSK3 reveals a high degree of homogeneity with a calculated molecular weight equal to the expected value of 52 kDa. (B) Stability dependence of the PfGSK3 melting temperature measured by nano differential scanning fluorimetry (nDSF) in the presence of different concentrations of ATP or dATP. Melting temperature increases with the concentration of ATP or dATP, indicative for binding ATP and its analogs. (C) Melting temperature of PfGSK3 measured by nDSF with or without 1 mM non-hydrolyzable ATP analogs. (D) Thermal unfolding profiles of PfGSK3 in presence of 500 mM sodium chloride or sodium sulfate show that PfGSK3 is preferentially stabilized by sulfate ions. (E) Stabilization of PfGSK3 by sulfate ions is concentration-dependent, indicating that stabilization effect is caused by specific sulfate ion binding. (F) Initial reaction velocity of GSK3 kinase reaction with 20 ng kinase, 6 µM ATP and substrate concentrations as indicated (GS-1 = peptide substrate of the human glycogen synthase). ATP consumption was quantified using the KinaseGlo assay (Promega) 5 and 10 min after start of kinase reaction. Lines were generated in GraphPad Prism using linear regression. Slope of each line represents reaction velocity (ATP consumption/min). Data points represent mean ± SD of three kinase reactions.

Supplementary figure 3

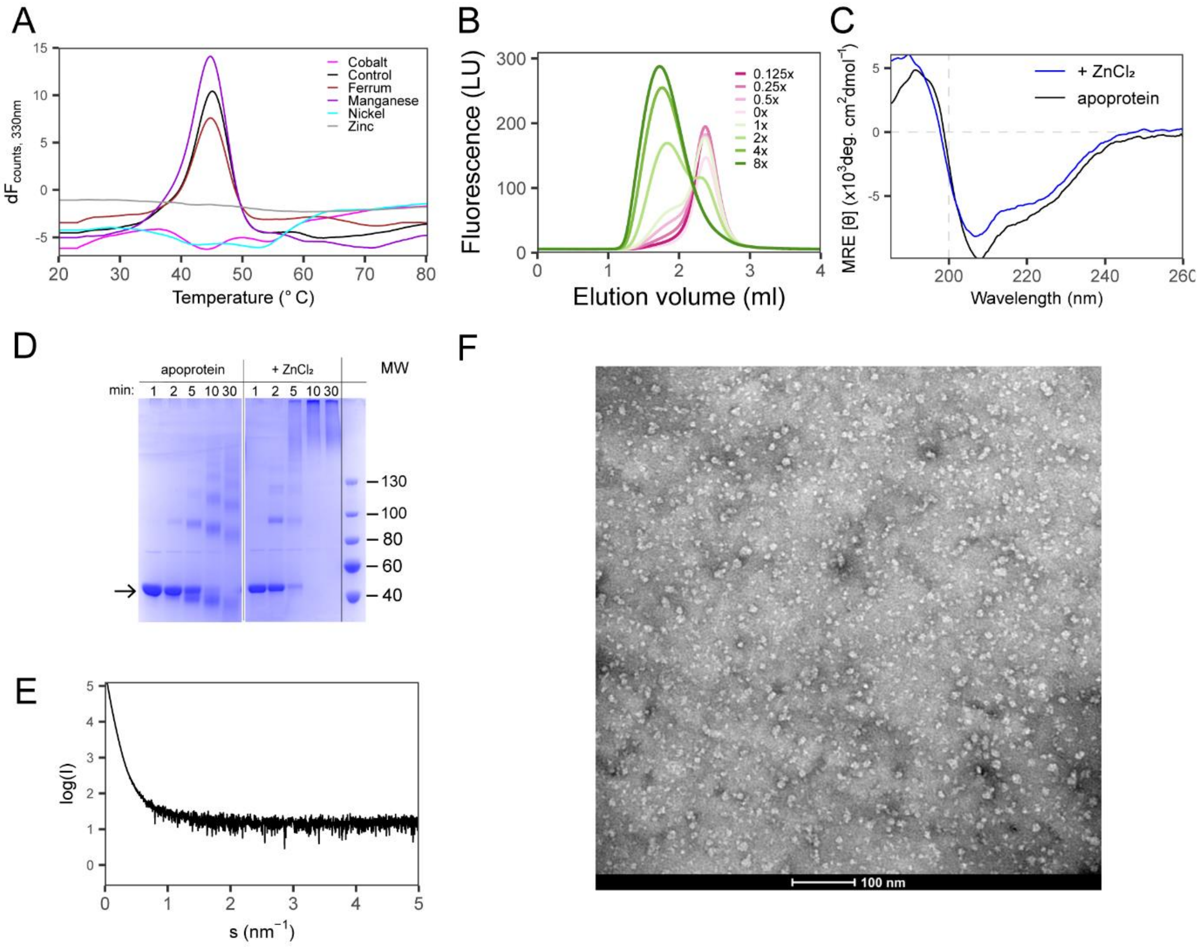


(A) Effect of heavy metals on thermal unfolding of PfGSK3. The unfolding trace was measured by nano differential scanning fluorimetry at a protein concentration of 1 mg/ml and a metal concentration of 1 mM. Presence of some heavy metal ions alters the unfolding profile of PfGSK3. (B) Effect of zinc chloride on the elution profile of PfGSK3 on a Superose 6 column. The experiment shows that the zinc ions induce the formation of high-MW particles in a concentration-dependent manner. The numbers indicate the molar access of metal over the protein. (C) Circular dichroism spectrum of PfGSK3 in the presence of zinc shows that the secondary structure is retained even in the induced high-MW particles. (D) Cross-linking experiment confirm that compared to apoprotein, PfGSK3 forms high-MW species that are induced by addition of zinc ions. (E) SAXS curve of PfGSK3 in the presence of zinc shows that the high-MW particles are large and heterogeneous. (F) Negative stain electron microscopy image of PfGSK3 high-MW particles induced by copper further stress the large sample heterogeneity.

Supplementary figure 4

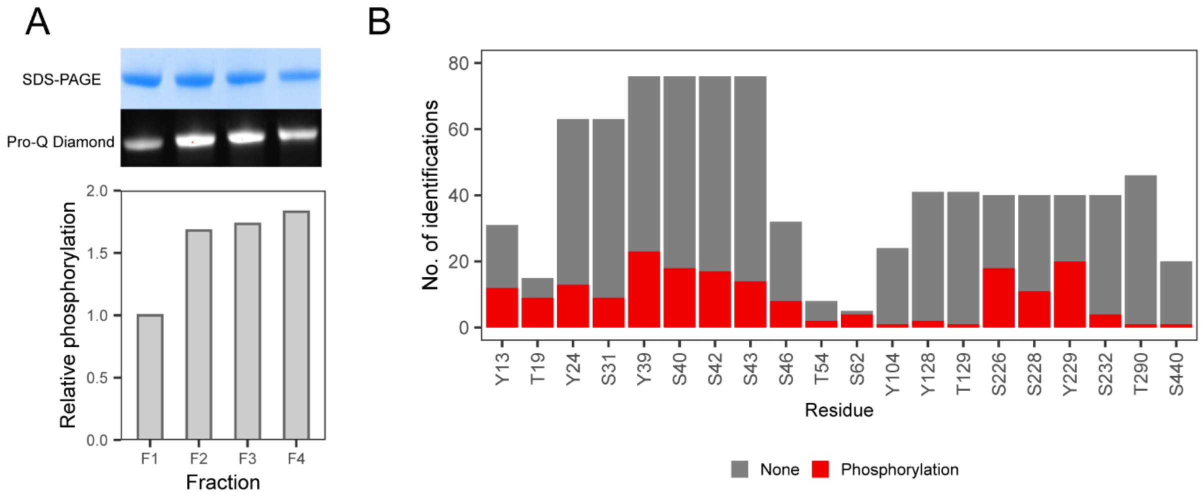


(A) The intensity ratio of bands between SDS-PAGE and Pro-Q diamond staining for individual IEX fraction shows that the later eluting species of PfGSK3 are more phosphorylated. The relative phosphorylation was calculated as ratio of relative band intensities with F1 band intensities as reference values. (B) LC MS-MS analysis of the PfGSK3 phosphorylation sites reveals heterogeneous phosphorylation of PfGSK3. All residues identified to be phosphorylated are shown.

Supplementary Figure 5

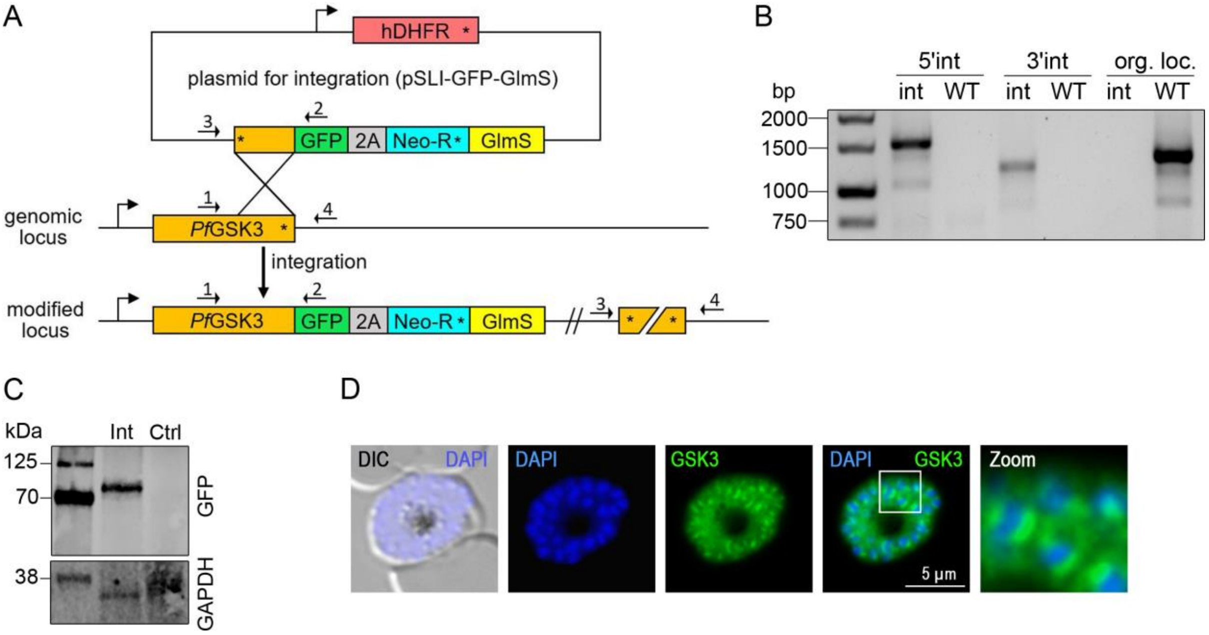


(A) Schematic representation of Selection Linked Integration (SLI) (84) used for C-terminal GFP-tagging. Introduction of a glms ribozym sequence in the 3'UTR allows conditional knock down upon addition of glucosamine (85, 110). Binding sites of primers are indicated by black arrows. hDHFR = human dihydrofolate reductase; GFP = green fluorescent protein; 2A = skip peptide; Neo-R = G418/neomycin-resistance gene; glmS = Glucosamine-activated ribozyme; angled arrows = promoters; asterisks = stop codon. (B) Integration PCR with genomic DNA isolated from PfGSK3-GFP-glms (int) or wild type parasites (WT). Primer combinations: 5'int (1+2); 3'int (3+4); org.loc. (1+4). Localization of primers as indicated in (A). Expected PCR products: 1+2 = 1470 bp; 3+4 = 1234 bp; 1+4 = 1344 bp. (C) Westernblot of PfGSK3-GFP-glms (Int) & wild type parasites (Ctrl) using mouse-anti-GFP. Expected size of PfGSK3-GFP: 80 kDa. GAPDH (111) served as a loading control.

Supplementary figure 6

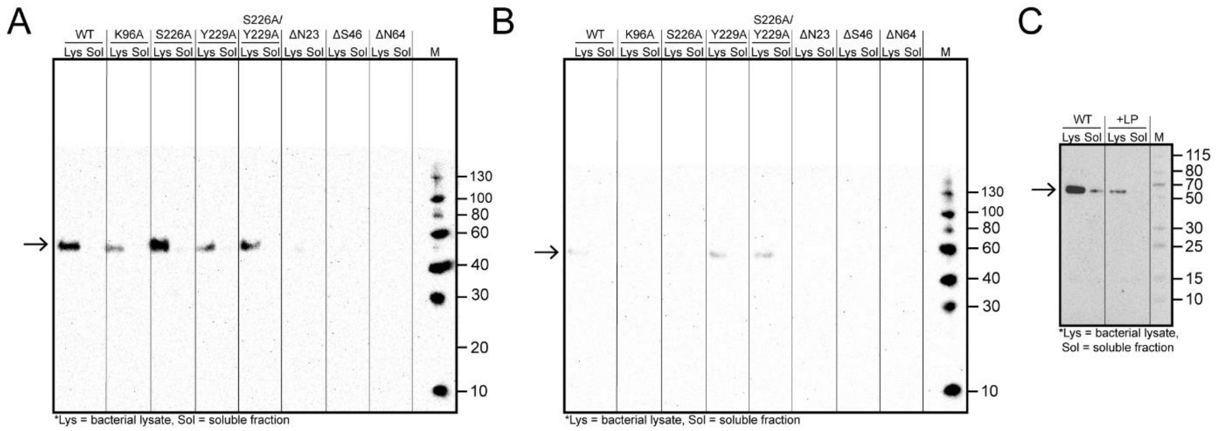


(A-B) Western blot analysis of expression test of PfGSK3 mutants in its ATP binding site (K96) or activation loop (S226 and Y229) and of PfGSK3 N-terminally truncated constructs at 25°C (A) and 18°C (B). The results show that the lower temperatures decrease the expression level and solubility of the proteins. (C) Western blot of test of PfGSK3 co-expression with λ-phosphatase shows that PfGSK3 dephosphorylation renders the protein insoluble.

Supplementary figure 7

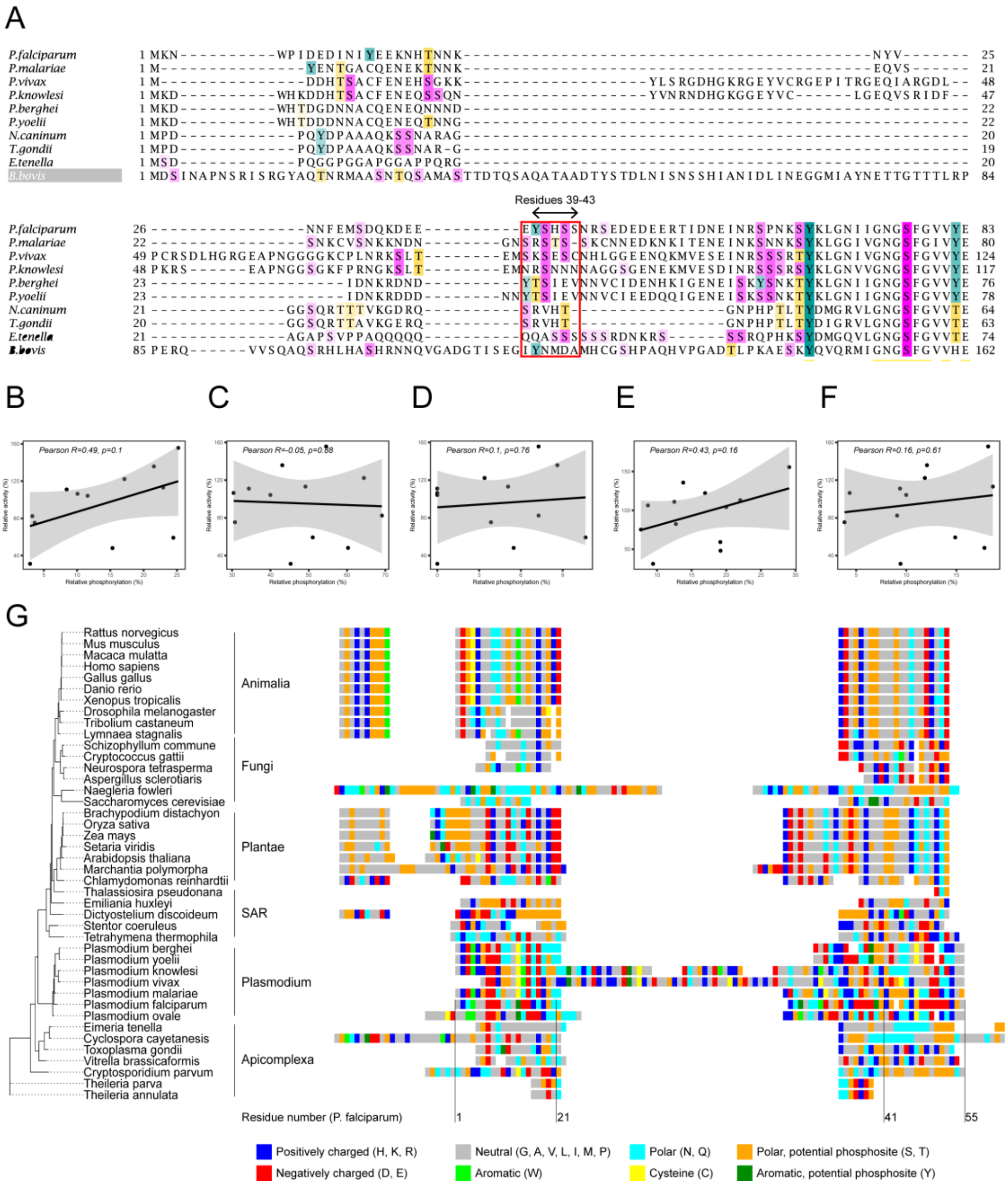


(A) Sequence alignment of PfGSK3 with homologous proteins from other *Plasmodia* and *Apicomplexa*. Only N-terminal residues are shown. Serine, threonine and tyrosine residues are depicted with magenta, yellow and cyan background, respectively. The region around regulatory N-terminal residues is highlighted in a red box. The alignment shows that residue S40 and neighboring residues are conserved in *Plasmodium* but not within *Apicomplexa*. (B-F) Correlation between the protein activity and the phosphorylation of residues Y39, S226, S228, Y229 and Y232, respectively. (G) Phylogenetic analysis of GSK3 shows that the N-terminus is unique in *Plasmodium* species as they separate into a clade distinct from other apicomplexans. Abbreviation SAR refers to a super group that includes Stramenopiles, Alveolates and Rhizaria.

Supplementary figure 8

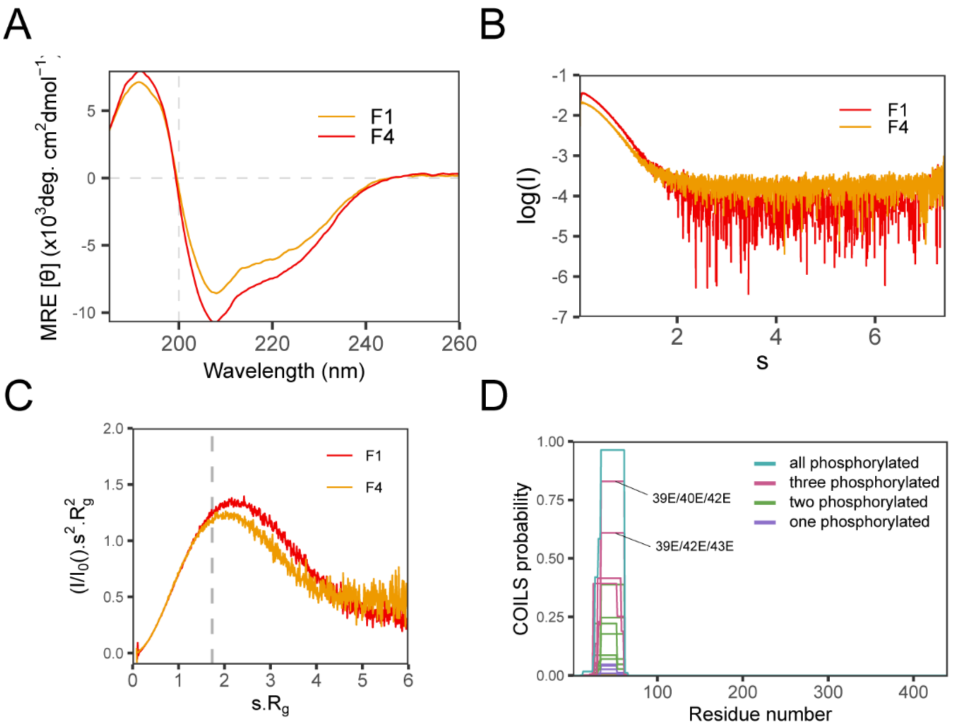


(A) Circular dichroism data measured on PfGSK3 fractions F1 and F4 indicates that the secondary structure does not change upon phosphorylation. (B) Small angle X-ray scattering data measured on PfGSK3 fractions F1 and F4. (C) Dimensionless Kratky plot representation of the SAXS data measured on fractions F1 and F4 shows that the phosphorylation of PfGSK3 (F4) leads to a less extended structural conformation. (D) *In silico* prediction of coiled-coil formation using the COILS server with PfGSK3 sequences with N-terminal residues (Y39, S40, S42 and S43) mutated to phosphomimetic glutamate residues used as an input.
